# CD4^+^ T follicular helper (Tfh) cells in human tonsil and blood are clonally convergent, but divergent from non-Tfh CD4^+^ cells

**DOI:** 10.1101/743187

**Authors:** Elena Brenna, Alexey N Davydov, Kristin Ladell, James E McLaren, Paolo Bonaiuti, Maria Metsger, Sarah C Gilbert, Teresa Lambe, David A Price, Suzanne L Campion, Dmitriy M Chudakov, Persephone Borrow, Andrew J McMichael

## Abstract

T follicular helper (Tfh) cells are fundamental for B cell selection and antibody maturation in germinal centers. Circulating Tfh (cTfh) cells constitute a minor proportion of the CD4^+^ T cells in peripheral blood, but their clonotypic relationship to Tfh populations resident in lymph nodes and the extent to which they differ from non-Tfh CD4^+^ cells has been unclear. Using donor-matched blood and tonsil samples we investigated T cell receptor (TCR) sharing between tonsillar Tfh cells and peripheral Tfh and non-Tfh cell populations. TCR transcript sequencing revealed considerable clonal overlap between peripheral and tonsillar Tfh cell subsets as well as a clear distinction between Tfh and non-Tfh cells. Furthermore, influenza-specific cTfh cell clones derived from blood could be found in the repertoire of tonsillar Tfh cells. Therefore, human blood samples can be used to gain insight into the specificity of Tfh responses occurring in lymphoid tissues, provided cTfh subsets are studied.

## INTRODUCTION

T follicular helper (Tfh) cells are specialized CD4^+^ T cells primarily found in germinal centers (GCs) in secondary lymphoid organs (Breitfeld et al., 2000; Kim et al., 2001; Schaerli et al., 2000). Tfh cells play a critical role in supporting B cell responses and selection of affinity matured antibodies (Breitfeld et al., 2000; Bryant et al., 2007; Ma et al., 2009). They mediate their effects via receptor-ligand interactions with B cells and production of cytokines such as IL-21, IL-4 and BAFF, which induce survival and proliferation in B cells and support antibody class-switching (Avery et al., 2008; Casamayor-Palleja et al., 1995; Liu et al., 1989). Expression of the chemokine receptor CXCR5 is fundamental for the migration of pre-Tfh cells to the T-B cell border in lymphoid tissues and the maturation of Tfh cells into B cell follicles and GCs along the follicular CXCL13 gradient (Ansel et al., 2000; Forster et al., 1996). In addition to CXCR5, Tfh cells also express PD-1 and ICOS (Choi et al., 2011; Dorfman et al., 2006; Haynes et al., 2007; Xu et al., 2013). Some memory CD4^+^ T cells in secondary lymphoid organs express intermediate levels of these markers, but Tfh cells within the GC (Tfh GC) express high levels of CXCR5 and PD-1, therefore a CXCR5^hi^PD-1^hi^ phenotype is commonly used to distinguish Tfh GC cells (Shi et al., 2018). Differences in expression of these surface markers reflect not only the location of CD4^+^ T cell sub-populations, but also their activation, differentiation and functional status (Crotty, 2018).

Populations of CD4^+^ memory T cells with similar characteristics to lymphoid Tfh cells have been described in the peripheral blood and are thought to represent circulating memory Tfh (cTfh) cells (Crotty, 2018; Hale and Ahmed, 2015). These peripheral cTfh cells express CXCR5 together with PD-1 and ICOS, but although a minute population of PD-1^hi^CXCR5^hi^ CD4^+^ T cells can be detected in peripheral blood, the majority cTfh population expresses these markers at much lower levels than Tfh GC cells (He et al., 2013; Vinuesa et al., 2016). Although there is some controversy about phenotypic definition of cTfh cells, it is accepted that CXCR5^+^CD4^+^ T cells promote Ig class-switching and plasmablast formation in co-culture with naïve or memory B cells (Bentebibel et al., 2013; He et al., 2013; Locci et al., 2016; Morita et al., 2011). Different subsets of cTfh cells have been distinguished: Th1-like (CXCR3^+^CCR6^−^), Th2-like (CXCR3^−^CCR6^−^) and Th17-like (CXCR3^−^CCR6^+^) cTfh cells, based on similarities with canonical Th CD4^+^ cell subpopulations (Bentebibel et al., 2013; Morita et al., 2011). The diversity of cTfh cells is also evidenced by the differences in cytokine production and transcription factor expression observed when cTfh cell subsets are co-cultured with naïve B cells in the presence of staphylococcal enterotoxin B (SEB). Th1-like subsets produce IFN-γ, Th2-like IL-4, IL-5 and IL-13 and Th17-like IL-17A and IL-22 (Bentebibel et al., 2013; Morita et al., 2011).

The Th2- and Th17-like subsets of cTfh cells provide better B cell help *in vitro* than Th1-like cTfh cells (Boswell et al., 2014; Locci et al., 2013; Morita et al., 2011), and the transcriptional profile of CXCR3^−^ cTfh shares strong similarity with that of Tfh GC cells (Locci et al., 2013). Nevertheless, during influenza virus infection in humans, where the CD4^+^ T cell response is highly Th1-biased, Th1-like (CXCR3^+^) cTfh cells help B cells produce virus-specific antibodies (Bentebibel et al., 2013; Pallikkuth et al., 2012). However, stimulation of Th1-like Tfh effector cells following infection or vaccination was associated with an inferior GC response and suboptimal antibody production (Bowyer et al., 2018; Cubas et al., 2015; Obeng-Adjei et al., 2015; Ryg-Cornejo et al., 2016).

Difficulties in sampling secondary lymphoid organs in humans have hampered direct comparison of lymphoid and peripheral Tfh cell populations. Furthermore, the ontogeny of memory cTfh cells and their relationship with effector Tfh GC cells are still poorly understood. Although a few previous studies have compared the phenotype (Chevalier et al., 2011; He et al., 2013) and transcriptional profile (Heit et al., 2017; Locci et al., 2013) of cTfh cell populations with Tfh GC cells, the extent of clonal convergence between these populations has not been fully explored. Phenotype and function are plastic features of CD4^+^ T cells, influenced by environmental stimuli, and therefore the same cell type can show a different phenotypic profile in different compartments or cell culture conditions (Sallusto et al., 2018). It has been unclear whether cTfh cells originate from CXCR5^−^ memory CD4^+^ T cells, from lymphoid tissue pre-Tfh or Tfh GC cells, or whether they comprise a distinct population. Understanding the relationship between Tfh and non-Tfh cell populations in the blood and lymphoid tissues is very important to inform interpretation of human studies, which typically investigate only the peripheral blood, because of the practical, ethical and consensual barriers to accessing lymphoid tissues.

The T cell receptor (TCR) is a fixed marker of clonotype identity in T cells. In this study, donor-matched tonsil and peripheral blood were collected from adult volunteers undergoing tonsillectomy and the clonotypic repertoire of Tfh and non-Tfh cell subsets was determined. TCR repertoire analyses revealed considerable overlap between peripheral and tonsillar Tfh cells, and a complete disparity between Tfh cells and CXCR5^−^ memory CD4^+^ T cells. In addition, memory Tfh and non-Tfh cells from blood reactive with the haemagglutinin (HA) protein from influenza virus, one of the most frequent seasonal infections, were studied (Alam and Sant, 2011; Fazilleau et al., 2007; Sant et al., 2018). HA-specific CD4^+^ memory T cells were enriched in CXCR3^+^ cTfh cells and HA-specific cTfh cell clones could be found in the tonsillar CXCR3^+^ Tfh population. These data confirmed the clonotypic overlap between peripheral and tonsillar Tfh cell populations in the context of antigen-specific memory CD4^+^ T cells. These results show that human blood samples, which are readily accessible, can be used to gain insight into the specificity of Tfh cell responses occurring in GCs, but only if cTfh populations are analyzed.

## RESULTS

### Identification of tonsillar and circulating Tfh populations in donor-matched samples

Matched peripheral blood and tonsil samples from adult donors were studied to directly compare cTfh and tonsillar Tfh populations. Multi-parameter flow cytometry-based characterization of CD4^+^ memory T cell populations in peripheral blood mononuclear cells (PBMCs) and mononuclear cells from tonsil (MNC-T) was performed (**Figure 1**). Expression of the Tfh markers CXCR5, PD-1, ICOS and the chemokine receptor CXCR3 on CD4^+^CD45RA^−^ memory T cells (identified as shown in **Figure S1A)** was evaluated (**Figure 1A**). A population with high expression of PD-1 and CXCR5 was observed only in tonsil and was not present in appreciable numbers in the peripheral blood samples. CXCR3 and ICOS were also differently expressed in blood and tonsil CD4^+^ memory T cells (**Figure 1B**). In tonsil, populations with lower expression of CXCR5 and PD-1 had the highest proportion of CXCR3^+^ cells, while in blood no significant difference was observed between CXCR5^+/–^ and PD-1^+/–^ subsets. By contrast, ICOS was maximally expressed in tonsil and blood populations with high expression of CXCR5 and PD-1, and it decreased as these two markers declined.

**Figure 1.**
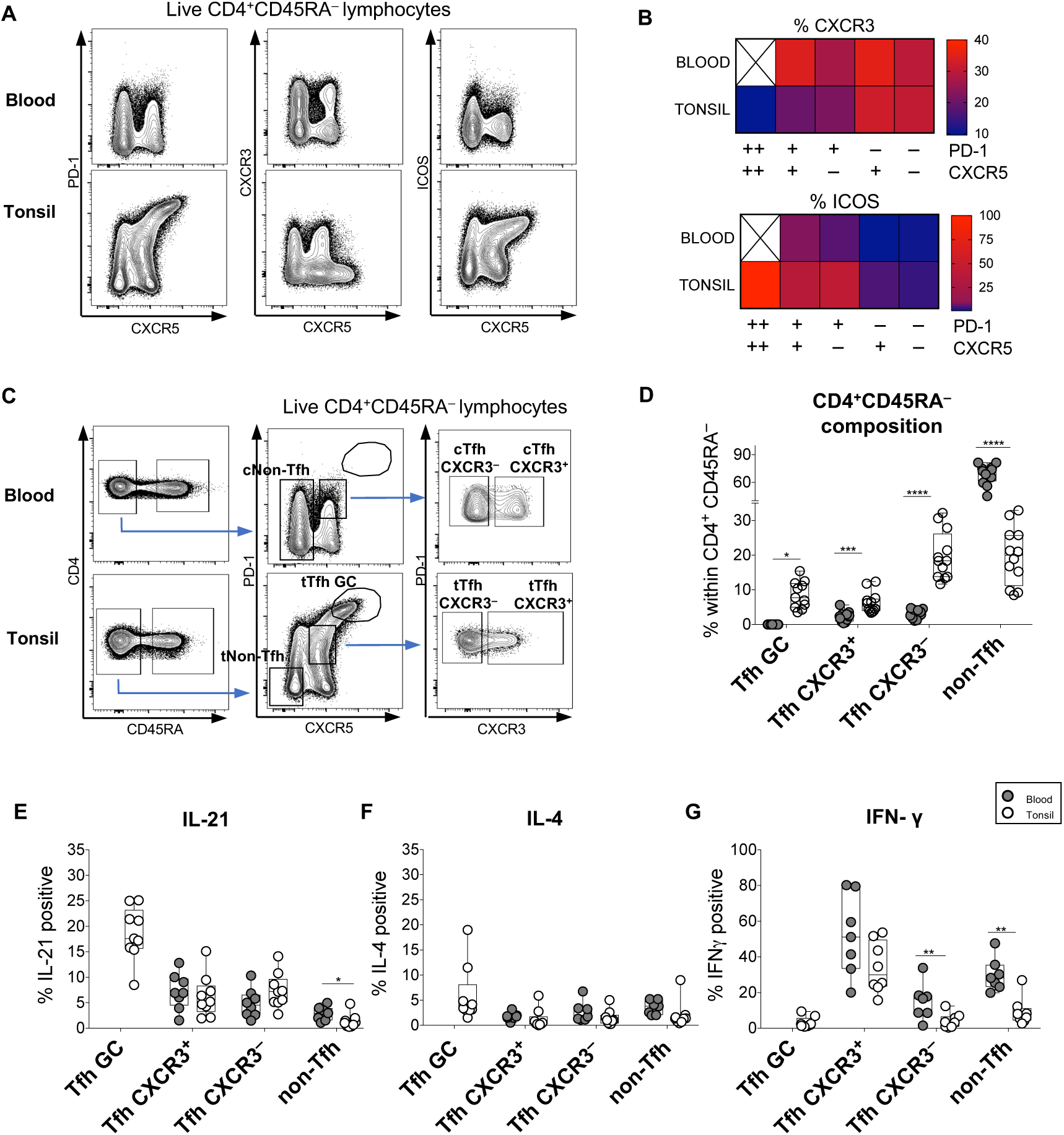
Identification of tonsillar and circulating Tfh populations in donor-matched samples. Matched PBMC and MNC-T from adult donors were stained *ex vivo*. Dot plot (**A**) illustrating the expression of CXCR5, PD-1, CXCR3 and ICOS within the total CD4^+^CD45RA^−^ cells in a representative matched blood and tonsil sample pair. Heat maps (**B**) showing mean % of CXCR3^+^ and ICOS^+^ cells within populations expressing different combinations of CXCR5 and PD-1 in 12 donors. The white crossed boxes indicate the near absence of the PD-1^hi^CXCR5^hi^ population in the peripheral blood. Dot plot (**C**) illustrating the gating strategy used to define populations of circulating CXCR5^+^PD-1^+^ Tfh (cTfh CXCR3^+^, cTfh CXCR3^−^) and CXCR5^−^ non-Tfh memory cells (cNon-Tfh); and tonsillar CXCR5^hi^PD-1^hi^ germinal centre Tfh (tTfh GC), CXCR5^int^PD-1^int^ (tTfh CXCR3^+^, tTfh CXCR3^−^) Tfh and CXCR5^−^ PD-1^−^ non-Tfh memory cells (tNon-Tfh) within the total CD4^+^CD45RA^−^ T cell pool in one representative donor. (**D**) shows the percentage of each Tfh/non-Tfh subset within the CD4^+^CD45RA^−^ memory T cell population in blood and tonsil from the 12 donors. Production of IL-21 (**E**), IL-4 (**F**), and IFN-γ (**G**) after PMA/ionomycin stimulation by Tfh and non-Tfh populations in 10 donors.

Subsets of CD4^+^ memory T cells were chosen for comparison based on the phenotypic observations made here and in prior studies (Boswell et al., 2014; Morita et al., 2011) (**Figure 1C**). Within the CD4^+^CD45RA^−^ population, cTfh CXCR3^+^ and cTfh CXCR3^−^ subsets of CXCR5^+^PD-1^+^ cells and cNon-Tfh cells (CXCR5^−^) were identified in peripheral blood. In tonsil, tTfh GC were identified as PD-1^hi^CXCR5^hi^, in contrast to the intermediate level PD-1^int^CXCR5^int^ population, in which tTfh CXCR3^+^ and tTfh CXCR3^−^ cells were characterized, and tNon-Tfh (CXCR5^−^PD-1^−^) cells were also studied. Gates were set based on the expression of PD-1, CXCR5 and CXCR3 in CD4^+^CD45RA^+^ naïve-enriched T cells (**Figure S1B**). The percentage of each population within CD4^+^CD45RA^−^ T cells was evaluated in samples collected from 12 donors (**Figure 1D**). The Tfh GC subset was not found in appreciable numbers in peripheral blood and Tfh subsets were less frequent in the peripheral blood than in tonsils, whilst non-Tfh cells were more abundant.

To define the functionality of phenotypically-distinct Tfh cell subsets, cytokine production from each subset was evaluated *ex vivo* after PMA/ionomycin stimulation, by flow cytometry. Data from 10 donors showed that Tfh GC cells were the major producers of IL-21 (**Figure 1E**) and IL-4 (**Figure 1F**), both crucial cytokines for Tfh cell differentiation and B cell survival (Nurieva et al., 2008; Reinhardt et al., 2009; Yusuf et al., 2010). The production of IL-21 was lower in the other Tfh cell subsets, and minimal in the non-Tfh cells (**Figure 1E**). No significant differences in IL-4 production were observed between Tfh CXCR3^+^, Tfh CXCR3^−^ and non-Tfh cells from both tonsil and blood (**Figure 1F**). In line with the low expression of CXCR3 in Tfh GC cells, this population also exhibited minimal production of IFN-γ (**Figure 1G**), consistent with published data (Ma et al., 2009; Ploquin et al., 2011). In contrast, the major producers of IFN-γ were Tfh CXCR3^+^ subsets from both tissues along with peripheral non-Tfh cells, which include the canonical Th1 phenotype cells known to be the highest producers of anti-viral cytokines within CD4^+^ cell subsets (Djuretic et al., 2007; Morita et al., 2011). Examples of cytokine staining of each cell subset are illustrated (**Figure S1C-F**).

### Clonotypic overlap between peripheral and tonsillar Tfh cell populations

To gain insight into the relationship between cTfh and tTfh populations, TCR repertoire analysis was performed. TCR clonotype sequence is a stable invariable characteristic of a T cell clone which does not change with cellular location or activation state and thus identifies clonal populations of T cells in different tissues. In order to compare the TCR repertoire of CD4^+^ memory T cell populations, frozen PBMCs and MNC-T from 4 matched donors were stained and tTfh GC, tTfh CXCR3^+^, tTfh CXCR3^−^ and tNon-Tfh from tonsil, and cTfh CXCR3^+^, cTfh CXCR3^−^ and cNon-Tfh from blood were separated by FACS (8,000-123,000 cells/subset). RNA was extracted and 5’RACE UMI-controlled sequencing of the CDR3 region of the TCR Vβ chain repertoire was performed.

The network analysis by Cytoscape in **Figure 2** illustrates the clonotypes shared between each population of blood CD4^+^ T cells: cTfh CXCR3^−^ (**Figure 2A**), cTfh CXCR3^+^ (**Figure 2C**) and cNon-Tfh (**Figure 2E**), and the different populations of tonsil (tTfh GC, tTfh CXCR3^+^, tTfh CXCR3^−^ and tNon-Tfh) cells, without showing the connections between the tonsil populations. The cTfh CXCR3^−^ population showed the greatest clonotype sharing with tTfh GC: a total of 6.45% of clonotypes were found to be shared between these populations overall. This contrasts with findings from a recent study where no clonal overlap was observed between tTfh GC and the blood cTfh populations studied here (Hill et al., 2019). 3.25% of clonotypes were uniquely shared between cTfh CXCR3^−^ and tTfh GC, whilst others were also co-shared with tTfh CXCR3^−^ subset, with which cTfh CXCR3^−^ also showed clonotype sharing: 2.05% of clonotypes were shared between all 3 of these populations, whilst a further 2.15% of clonotypes were uniquely shared between the cTfh CXCR3^−^ and tTfh CXCR3^−^ subsets (**Figure 2A**). By contrast, cTfh CXCR3^+^ and cNon-Tfh populations each showed extensive repertoire overlap only with the tonsil CD4^+^ population to which they were phenotypically most closely related: 12.8% of clonotypes were uniquely shared between the cTfh CXCR3^+^ and tTfh CXCR3^+^ populations (**Figure 2C**), whilst cNon-Tfh cells uniquely shared 19.2% with tNon-Tfh cells (**Figure 2E**). The percentage of clonotypes shared between populations in each of the 4 donors analyzed is shown in **Figures 2B**, **2D** and **2F**, confirming the overlap between cTfh CXCR3^−^ and both tTfh GC and tTfh CXCR3^−^ populations (**Figure 2B**), the extensive clonotype sharing of cTfh CXCR3^+^ with tTfh CXCR3^+^ cells (**Figure 2D**) and of cNon-Tfh with tNon-Tfh cells (**Figure 2F**). The heat map (**Figure 2G**) displays the number of clonotypes shared between populations for all 4 donors.

**Figure 2.**
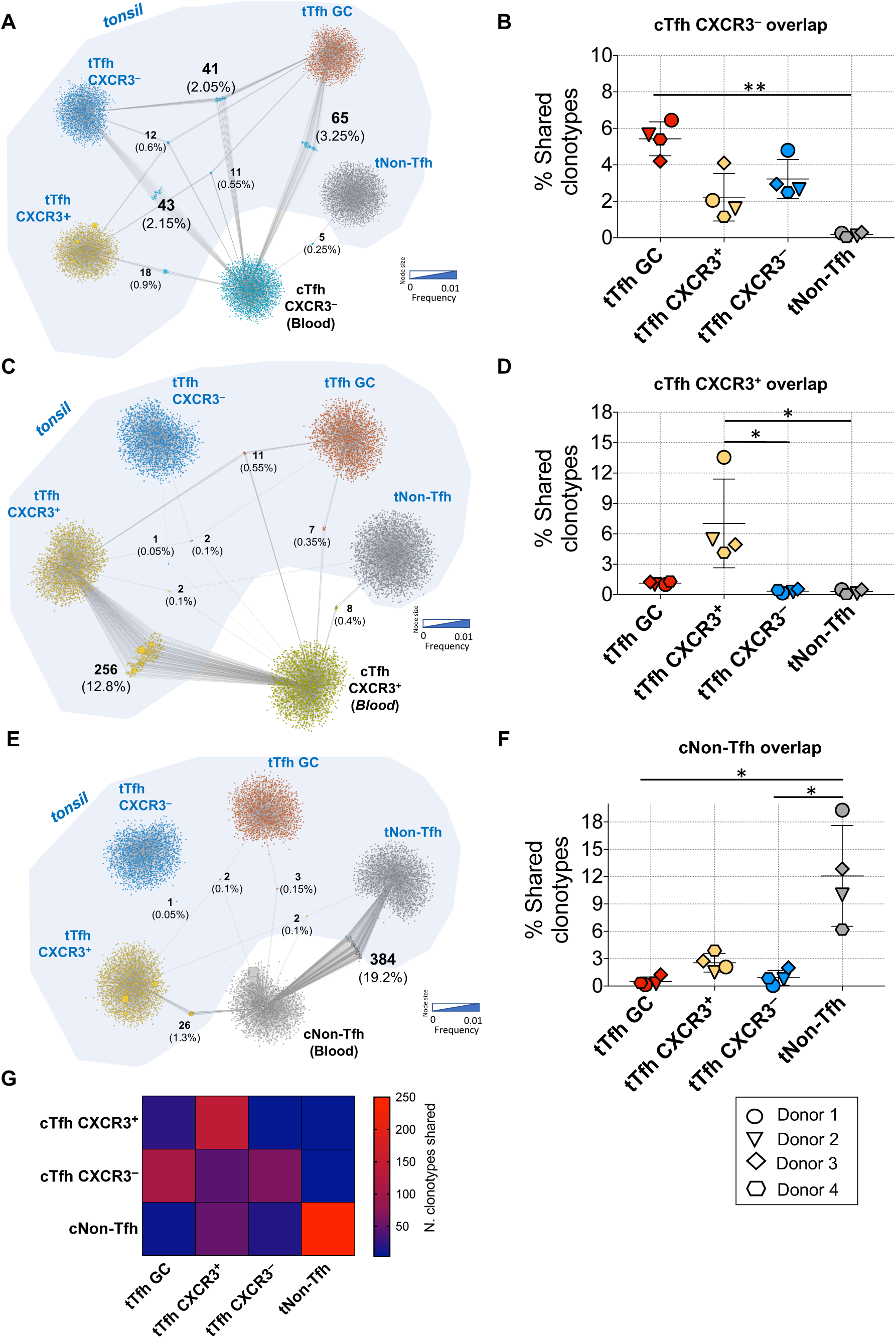
Clonotypic overlap between peripheral and tonsillar Tfh cell populations. The top2000 TCR Vβ CDR3 nucleotide sequences in each population were analysed using Cytoscape networking. As an example, clonotype sharing between peripheral blood cTfh CXCR3^−^ (**A**), cTfh CXCR3^+^ (**C**) and cNon-Tfh (**E**) and 4 tonsillar populations: tTfh CXCR3^+^ in yellow, tTfh CXCR3^−^ in blue, tTfh GC in red and tNon-Tfh in grey in one representative donor (donor 1) is illustrated. Each square represents a different clonotype; the size of the square is proportional to the clonotype’s normalized frequency within the parent population (for shared clonotypes the highest normalized frequency is shown). The number of clonotypes shared between different populations is shown in bold and the respective percentage is indicated below. To compare the overlap between blood cTfh CXCR3^−^ (**B**), cTfh CXCR3^+^ (**D**) and cNon-Tfh (**F**) subsets and tonsillar populations, the percentage of the top2000 most frequent clonotypes shared between populations was calculated in each of the 4 donors (mean ± SEM). A heat map (**G**) depicting the average number of clonotypes shared is also shown.

For a deeper TCR repertoire comparison, it was important also to consider the size of each clonotype (weighted analysis) to gauge the repertoire overlap in terms of cell numbers. For each tonsil and blood comparison the normalized F2 metric of VDJtools (Shugay et al., 2013) was calculated. The greater the F2 value between two subsets, the higher the number of T cells bearing the shared TCR variants (Izraelson et al., 2017). This analysis confirmed the repertoire similarity of cTfh CXCR3^−^ with tTfh GC and tTfh CXCR3^−^ populations, and identified a connection between cTfh CXCR3^−^ and tTfh CXCR3^+^ cells (**Figure 3A**). Likewise, the F2 metric also validated the overlap of cTfh CXCR3^+^ with tTfh CXCR3^+^ cells (**Figure 3B**) and the extensive overlap of cNon-Tfh with tNon-Tfh cells, as well as identifying weaker connections with tTfh populations (**Figure 3C**).

**Figure 3.**
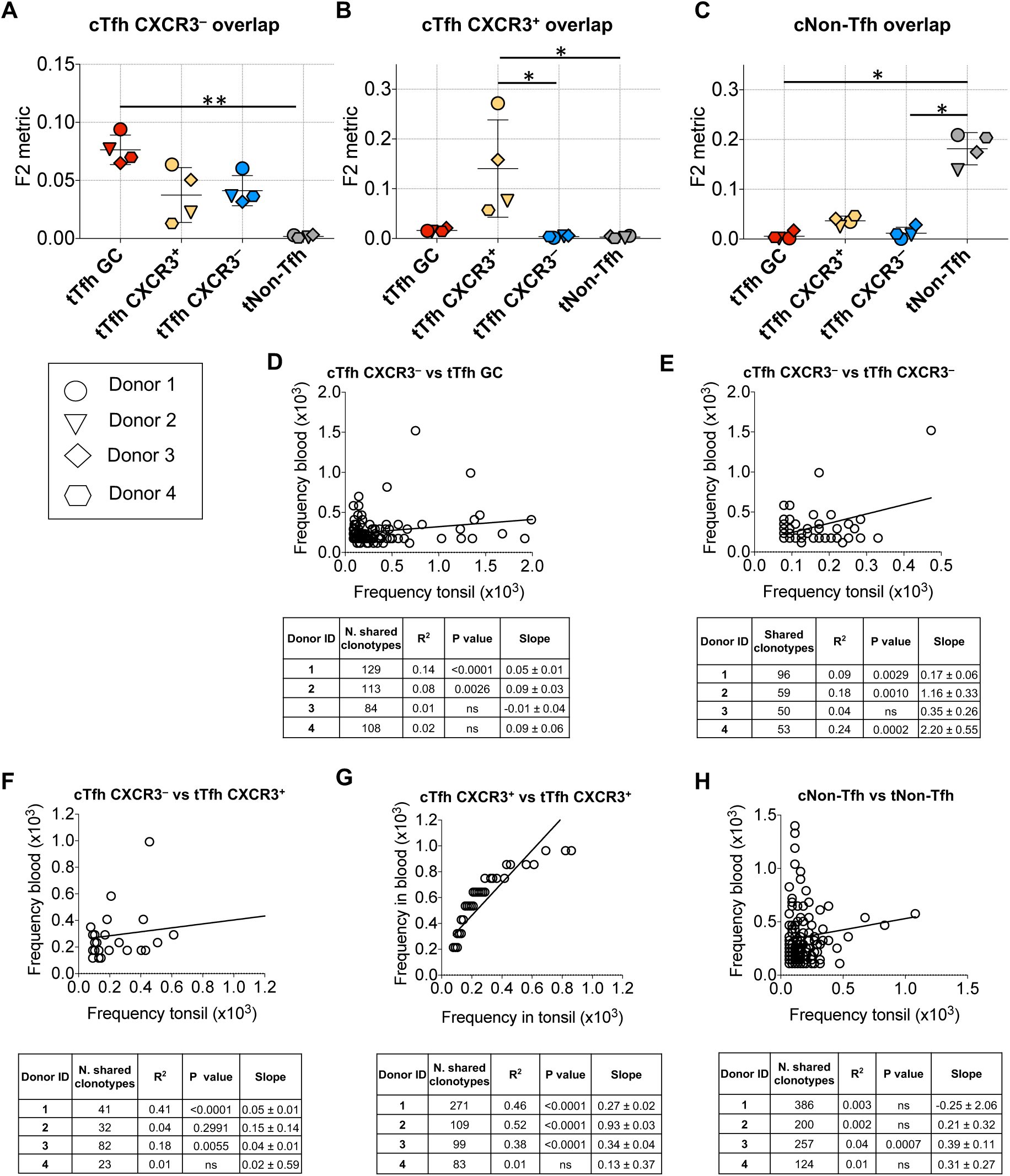
Size similarities in clonotypes shared between cTfh and tTfh. Overlap between TCR Vβ CDR3 nucleotide sequences of the top2000 most frequent clonotypes of blood cTfh CXCR3^−^ (**A**), cTfh CXCR3^+^ (**B**) and cNon-Tfh (**C**) populations with each tonsil population, as represented by the normalized F2 metric (mean ± SEM). Linear regression analysis of the correlation between the within-population frequency of the shared clonotypes in the two parent populations, where clonotype sharing between populations was observed: cTfh CXCR3^−^ with tTfh GC (**D**), tTfh CXCR3^−^ with cTfh CXCR3^−^ (**E**), cTfh CXCR3^−^ with tTfh CXCR3^+^ (**F**), cTfh CXCR3^+^ with tTfh CXCR3^+^ (**G**) and cNon-Tfh with tNon-Tfh (**H**). Each graph illustrates a representative donor and the tables show the number of shared clonotypes, the square of the Pearson correlation coefficient (R^2^), the slope value for each donor and the P value of the significance of the difference in the slope from zero (ns= not significant).

To visualize the similarity between the two compartments better, where clonotype sharing was observed the frequencies of each shared clonotype were analyzed by linear regression. Although there was consistent overlap between cTfh CXCR3^−^ and tTfh GC cells (**Figure 3D**), comparison of the relative sizes of clones shared by the two different compartments was significant in only 2 of the 4 donors, with a low R^2^ (0.08 and 0.14). In all cases, the slope of the correlation was <1, indicating that in tonsils most of the clones are more expanded than in the blood. A similar scenario was observed for cTfh CXCR3^−^ versus tTfh CXCR3^−^ subsets (**Figure 3E**) and cTfh CXCR3^−^ versus tTfh CXCR3^+^ populations (**Figure 3F**). The correspondence between cTfh CXCR3^+^ and tTfh CXCR3^+^ populations remained the strongest, as the sizes of the clonotypes shared between these populations were also very similar, significant (P<0.0001) in 3 of the 4 donors (**Figure 3G**). For the cNon-Tfh and tNon-Tfh comparison, despite the high number of clonotypes shared (124-386), the size of the clonotypes was very different between blood and tonsil, making the linear correlation significant in only 1 of the 4 donors (**Figure 3H**).

### Repertoire diversity correlates with the degree of overlap from *in silico* resampling analysis

To compare the repertoire diversity of each population the non-parametric index, Chao1 (**Figure 4A**), was calculated to estimate the richness of the number of unobserved elements in the set considered (Colwell et al., 2012; Eren et al., 2012). The normalized Shannon-Wiener index was also calculated to measure the entropy of each TCR sequence population (**Figure 4B**), based on the uncertainty of predicting the next sequence in a string. The two parameters describe the repertoire diversity from different perspectives: Chao1 is based on relative distributions of low frequency clonotypes, while Shannon-Wiener takes into account large clonal expansions (Izraelson et al., 2017). Notably, an increase of diversity was observed in blood from cTfh CXCR3^+^ to cTfh CXCR3^−^ and cNon-Tfh cells, likewise in tonsil tTfh GC cells were less diverse than tNon-Tfh subsets, which showed the highest value of both Chao1 and normalized Shannon-Wiener indices.

**Figure 4.**
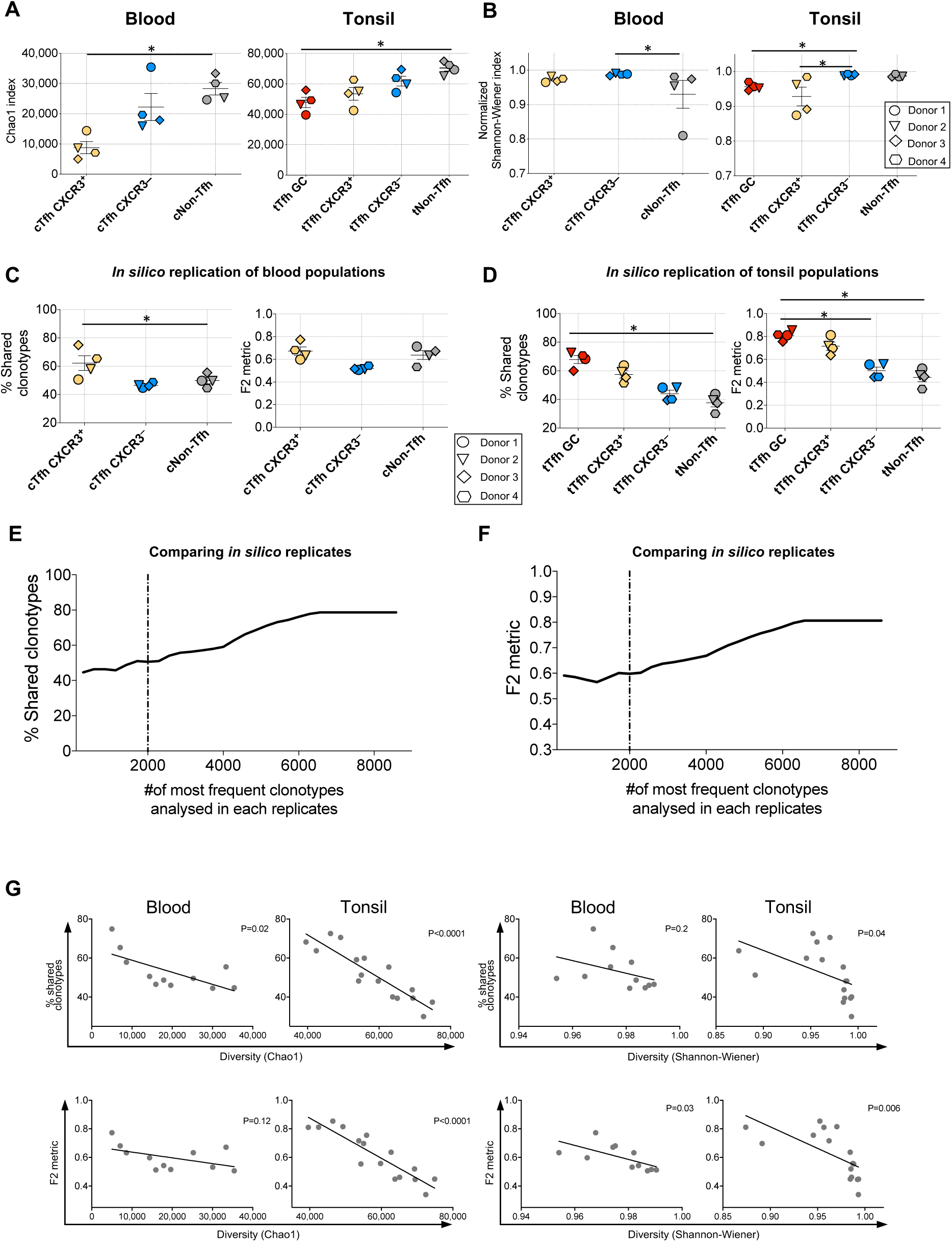
Repertoire diversity correlates with the degree of overlap from *in silico* resampling analysis. Chao1 index (**A**) and normalized Shannon-Wiener index of the top2000 frequencies (**B**) for each population of cells analysed in blood and tonsil in the 4 donors (mean ± SEM). Clonotypes of each subset were resampled using the bootstrap method. The percentage of the top2000 shared clonotypes and the normalized F2 metric (mean ± SEM) of the overlap between the two replicates of blood (**C**) cTfh CXCR3^+^, cTfh CXCR3^−^ and cNon-Tfh as well as tonsil (**D**) tTfh GC, tTfh CXCR3^+^, tTfh CXCR3^−^ and tNon-Tfh populations is shown for all the 4 donors (mean ± SEM). Unweighted (**E**) and weighted (**F**) analysis of the replicate overlap at increasing number of clonotypes considered from a representative population (cTfh CXCR3^+^ from donor 1). The graphs (**G**) show the linear regression correlation between the repertoire overlap, expressed either as percentage of shared clonotypes or normalized F2 metric, with both the diversity indexes (Chao1 and normalized Shannon-Wiener) in all blood and tonsil populations from the 4 donors analysed.

In order to improve understanding of the magnitude of the overlap observed between CD4^+^ T cell subsets in blood and tonsil, which was based on analysis of a limited number of cells from each population, an *in silico* replication using the bootstrap method was applied to re-sample the clonotypes in each subset. When re-sampling the same subset twice, populations from the blood (**Figure 4C** and **S2A**) showed on average an overlap between sampling 1 and sampling 2 of 50%, with the cTfh CXCR3^+^ cells slightly higher (60-70%). In tonsil populations (**Figure 4D** and **S2A**), the highest reproducibility was reported for tTfh GC cells, with decreasing levels of overlap being observed for tTfh CXCR3^+^ and tTfh CXCR3^−^ populations and the lowest with tNon-Tfh cells. A similar trend was reported from the weighted analysis represented by the F2 metric analysis.

When increasing the number of clonotypes considered, the prediction did not reach the complete overlap, but rather a plateau of 60% of unweighted (**Figure 4E**) and 0.8 weighted (**Figure 4F**) analysis respectively. As expected, the degree of overlap was inversely related to population diversity (**Figure 4G**), and was more pronounced for tonsil compared to blood, confirming that when a population is less diverse, the degree of overlap on *in silico* resampling is higher. The overlap of the same population was also assessed experimentally in non-Tfh (**Figure S2B**) and Tfh cells (**Figure S2C**) from 3 additional donors, and was found to be <30% for non-Tfh and <52% for Tfh subsets.

These results show that even when re-sampling exactly the same set of sequences or sequencing two vials of the same sorted population, the repertoire overlap is never complete (100% and F2=1) and it changes depending on the diversity of the subset considered and on the size of the sample (Shugay et al., 2013). This implies that the degree of overlap observed experimentally is an underestimate, because of the relatively small sample size, reinforcing the conclusion that there is substantial repertoire overlap between blood and tonsil Tfh populations.

### Tfh cell subsets are clonally distinct from Non-Tfh cells

Whether Tfh cells make up a distinct population of CD4^+^ T cells with a unique program of differentiation or are a subset of the memory CD4^+^ T cell compartment in a transient activation state is still debated (Baumjohann et al., 2013; Crotty, 2014; Herati et al., 2017; Vinuesa et al., 2005; Weber et al., 2015).

Cytoscape networking analysis revealed only a very low level of TCR Vβ repertoire overlap between cTfh CXCR3^+^, cTfh CXCR3^−^ and cNon-Tfh populations in the blood compartment (**Figure 5A**). This was true for both comparisons of cTfh CXCR3^+^ versus cTfh CXCR3^−^ (0.8%) and cTfh subsets versus cNon-Tfh (0.3-1%) populations. In addition, very few clonotypes were shared between all 3 populations (0.2%). These results were consistent in all 4 donors analyzed (**Figure S3**). Similarly, tTfh cells (either CXCR3^+^, CXCR3^−^ or GC) were also notably distinct from the tNon-Tfh population (**Figure 5B**), with very little repertoire overlap (0.1-0.3%). In contrast, the 3 tTfh populations showed considerable overlap with each other: tTfh CXCR3^+^ with tTfh CXCR3^−^ and Tfh GC (0.8-3.1%), tTfh CXCR3^+^ with tTfh GC (3.6-7.4%), tTfh CXCR3^−^ with tTfh GC (4-14.9%), and tTfh CXCR3^+^ with tTfh CXCR3^−^ (2.7-6.3%) (**Figure 5C**). Although the extent of the repertoire overlap between the tTfh populations was variable in different donors, clonotype sharing between the 3 tTfh cell populations was observed in all 4 donors, especially between the tTfh CXCR3^−^ and tTfh GC subsets. Normalized F2 metric analysis confirmed the overlap between tTfh CXCR3^−^ and tTfh GC subsets and the repertoire distance between all tTfh populations and tNon-Tfh (**Figure 5D**). Despite the numerous clonotypes shared between tTfh CXCR3^+^ and tTfh CXCR3^−^ (70-172), tTfh CXCR3^+^ and tTfh GC (89-214), tTfh CXCR3^−^ and tTfh GC (96-360), the relative immunodominance of the shared clonotypes differed substantially: the correlations were R^2^<0.36 and in the majority of the cases not significant (**Figure 5E**).

**Figure 5.**
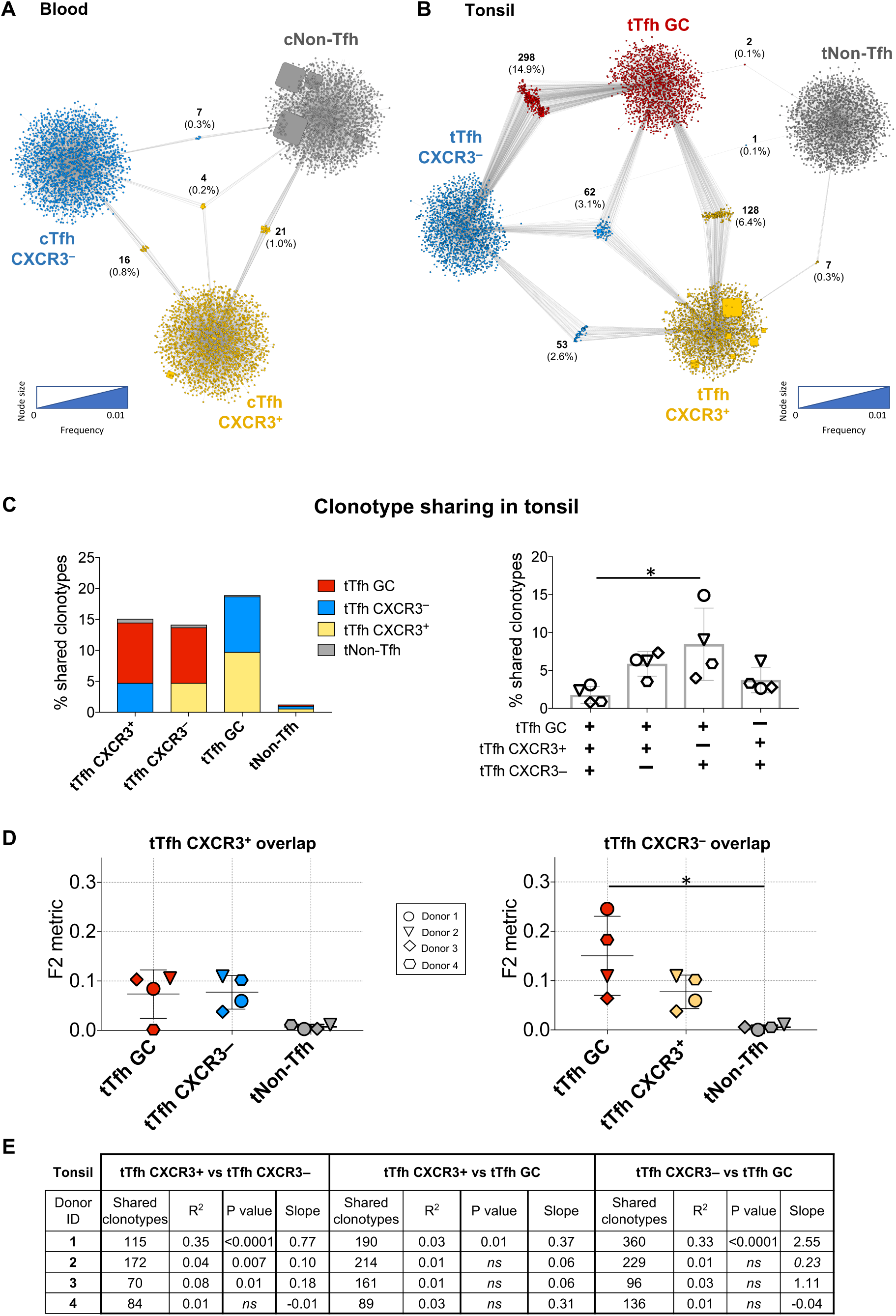
Tfh cell subsets are clonally distinct from Non-Tfh cells. The top2000 TCR Vβ CDR3 nucleotide sequences of each population were analysed by Cytoscape networking. Examples of (**A**) clonotype sharing between peripheral blood cTfh CXCR3^+^ (yellow), cTfh CXCR3^−^ (blue) and cNon-Tfh (grey) populations and (**B**) clonotype sharing between tonsillar populations tTfh CXCR3^+^ (yellow), tTfh CXCR3^−^ (blue), tTfh GC (red) and tNon-Tfh (grey) populations in one representative donor (donor 1) are illustrated. Results are expressed as in Figure 2. The percentage of the top2000 total shared clonotypes in addition to the uniquely shared clonotypes between tonsil populations was calculated in each of the 4 donors (mean ± SEM) (**C**). The normalized F2 metric of the top2000 frequencies was also calculated for the overlap between tTfh CXCR3^+^ and tTfh CXCR3^−^ subsets and each of the other populations analysed in the 4 donors (mean ± SEM (**D**). The table shows the number of shared clonotypes, the square of the Pearson correlation coefficient (R^2^) in their size within populations, the slope value and the P value of the difference in the slope from zero (ns= not significant) (**E**).

### Influenza virus HA-specific clones are enriched in CXCR3^+^ cTfh

To compare the repertoire of antigen-specific Tfh and non-Tfh cells in peripheral blood and their representation in tonsil populations, responses to the haemagglutinin (HA) protein H3 from influenza virus A/Wisconsin/67/2005 were studied (Alam and Sant, 2011; Fazilleau et al., 2007; Sant et al., 2018). To assess the relative abundance of memory CD4^+^ T cells reactive with HA protein the T cell library method (Geiger et al., 2009) was used to determine the precursor frequency of HA-reactive cells in isolated memory CD4^+^ T cell populations and to characterize the cTfh and cNon-Tfh CD4^+^ memory repertoire in 3 healthy donors (**Figure 6A**). All 3 donors showed enrichment of HA-specific T cells in the cTfh CXCR3^+^ and cNon-Tfh populations. Few responding HA-specific cTfh CXCR3^−^ cells were found, and they were only detected in donor A. The precursor frequency of antigen-specific cells per million was calculated within each subset (**Figure 6B**). Donors A and B showed the highest HA-specific precursor frequency in both cTfh CXCR3^+^ and cNon-Tfh populations (530-550 and 280-150 antigen-specific cells/million). The differences observed in the 3 donors analyzed likely reflects the diverse prior antigenic exposure of each individual.

**Figure 6.**
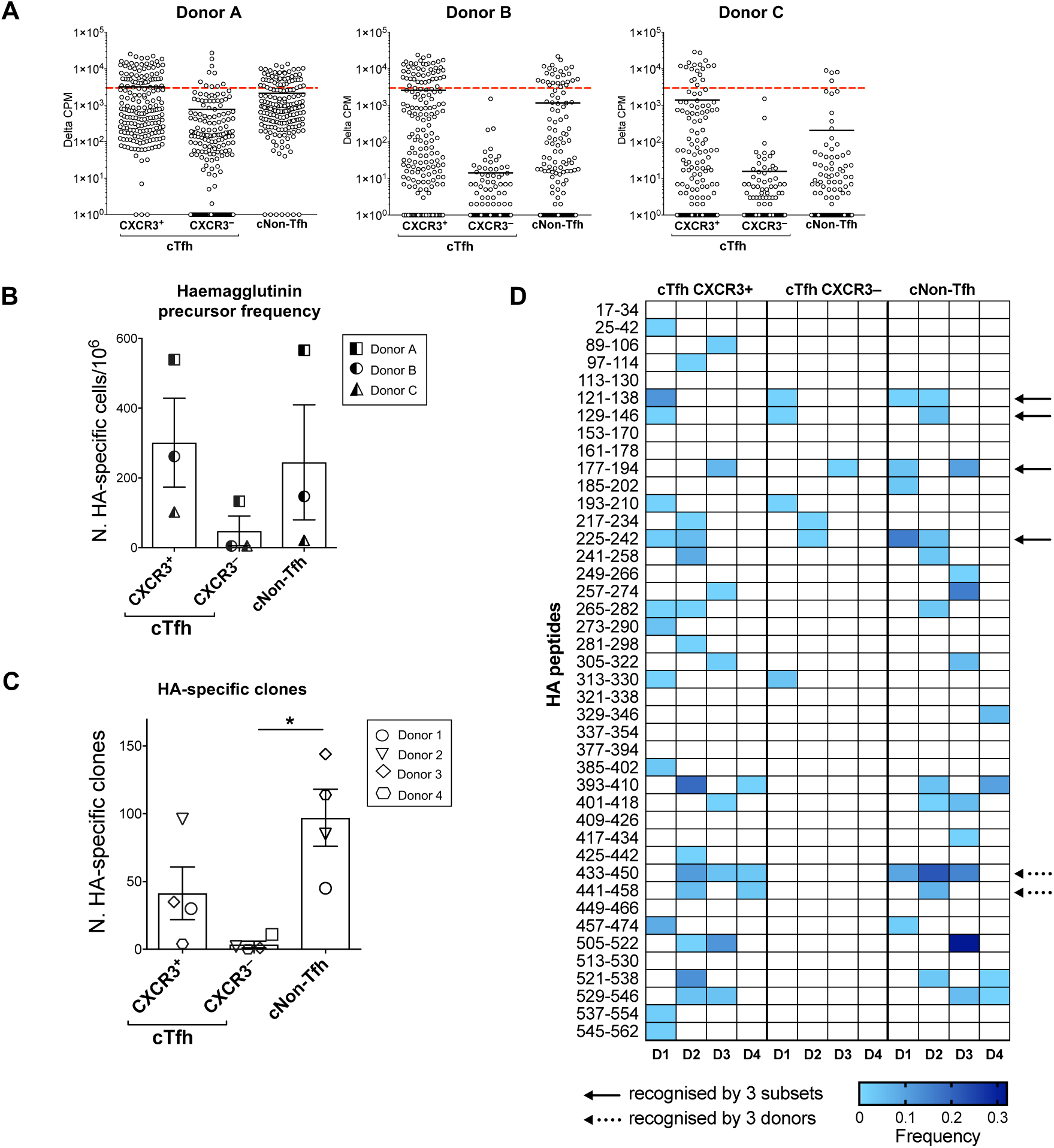
Influenza virus HA-specific clones are enriched in CXCR3^+^ cTfh. T cell libraries generated from Tfh and non-Tfh subsets sorted from 3 healthy blood donors were tested against the influenza H3/Wisconsin HA protein. The proliferation of each individual cell line (increase in counts above background) (Delta CPM) is shown (mean ± SEM). The red dotted line represents the cut-off used for identification of a positive response (defined as >3000 cpm, with a stimulation index >5) (**A**). Calculated precursor frequency of HA-specific cells per 10^6^ cells in each subset (mean ± SEM) (**B**). Bar graph showing the number of influenza H1/California HA-specific T cell clones generated from each cell subset in the 4 further donors from whom matched tonsil samples were available (**C**). Epitopes recognised by HA-specific cTfh CXCR3^+^, cTfh CXCR3^−^ and cNon-Tfh clones were mapped and the frequency of clones recognising each peptide, normalized to the total frequency of HA-specific clones in all donors (D1, D2, D3, D4) is plotted in the heat map (**D**). The solid arrows highlight the peptides recognized by 3 subsets, the dashed arrows the peptides recognized by 3 donors.

Having shown that HA-specific cTfh cells are present in the peripheral blood, the antigen-specific cTfh repertoires of 4 further donors, from whom tonsil samples were available in addition to peripheral blood samples (donors 1, 2, 3 and 4), were compared with the entire repertoire of T cells in tonsillar cell subsets from the same individual. To confirm influenza virus exposure, plasma samples from the 4 donors were screened for the presence of HA-specific antibodies, evaluating IgG reactivity with HA protein from influenza virus strain A/California/07/2009, H1N1 (**Figure S4A**) and IgG reactivity with trivalent (H1, H3 and B) influenza Vaccine 2017 (**Figure S4B**) by ELISA. IgG antibodies reactive with both HA protein and TIV were detected in all 4 donors. Tfh cell clones specific for HA protein (H1) were then generated from peripheral blood cTfh CXCR3^+^, cTfh CXCR3^−^ and cNon-Tfh populations of each donor. As shown in **Figure 6C**, the highest numbers of HA-specific clones were derived from either cTfh CXCR3^+^ or cNon-Tfh populations. These were the subsets in which antigen-specific cloning efficiency was greatest (**Figure S4D**). Based on the percentage of activated ICOS^+^CD25^+^ cells after 7 days of stimulation (**Figure S4C**) and the proportion of the clones generated from the activated populations that were subsequently confirmed to be HA-reactive by proliferation, it was possible to gain estimates of the number of HA-specific cells within each subset (**Figure S4E**), or within the CD4^+^ memory compartment (**Figure S4F**). The data confirmed that HA-specific T cells are enriched in the cTfh CXCR3^+^ and cNon-Tfh populations, while, as expected, only a few HA-specific cells were found in the cTfh CXCR3^−^ compartment (Bentebibel et al., 2013; Pallikkuth et al., 2012).

The epitopes recognized by HA-specific T cell clones were then mapped using a matrix of overlapping peptides from HA/California (H1). The data from cTfh CXCR3^+^ and cNon-Tfh (**Figure 6D**) clones highlighted the breath of HA epitopes recognized in each donor and suggested a difference in the pattern of epitopes recognized by cTfh and cNon-Tfh subsets, as only a few peptides per donor were recognized by all the three types of cells (solid arrows). Epitopes recognized by at least 3 donors are highlighted with a dashed arrow: peptide 433-450 was the most universally targeted, being recognized by cTfh CXCR3^+^ clones from donors 2, 3 and 4 and by cNon-Tfh clones from donors 1, 2 and 3. The number of cTfh CXCR3^−^ clones was low, and none of the clones recognized epitopes targeted in at least 3 donors, including 433-450 and 441-458.

### HA-specific cTfh CXCR3^+^ T cell clonotypes can be detected in the tTfh CXCR3^+^ subset

To understand whether clones with the same peptide specificity also shared the same TCR sequence, the CDR3α and CDR3β regions of a subset of HA-specific clones from donors 1, 2 and 3 were sequenced (**Figure 7A**). These data showed that when cTfh CXCR3^+^, cTfh CXCR3^−^ and cNon-Tfh cell clones were specific for the same peptide (for example, peptide 225-242 for donors 1 and 2 and peptide 177-194 for donor 3), they did not share the same CDR3 region; therefore, those cells belonged to different T cell clones. In some cases, even if 2 clones were specific for the same peptide and belonged to two different lineages, they used TCRα or TCRβ chains from the same family. For instance, in donor 2, the clones 49 (cTfh CXCR3^+^), 1 (cTfh CXCR3^−^) and 42 (cNon-Tfh) specific for peptide 225-242 exhibited different CDR3α sequences, but all three TCRα chains belonged to the same TRAV23/DV6 family. Also, in donor 3, clones 1 (cTfh CXCR3^+^) and 1 (cTfh-CXCR3^−^) specific for peptide 177-194, which again had different CDR3α sequences, possessed TCRβ chains belonging to the same family (TRBV2). Additional sequences from donor 1 are shown in **Figure S5A**.

**Figure 7.**
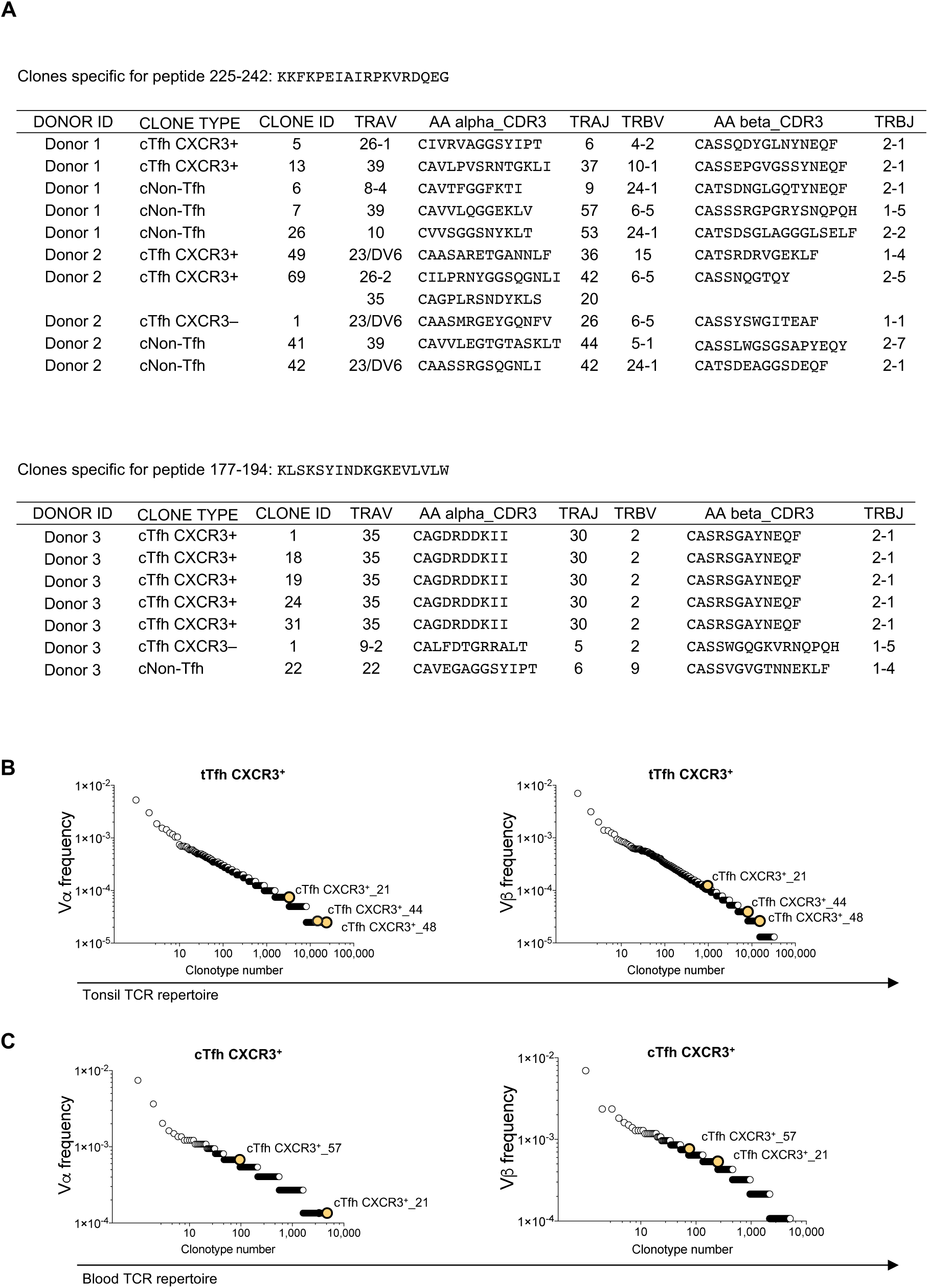
HA-specific cTfh and cNon-Tfh clones do not overlap and HA-specific cTfh CXCR3^+^ T cell clonotypes can be detected in the tTfh CXCR3^+^ subset. Vα and Vβ TCR CDR3 sequencing was performed on clones specific for the same HA H1/California peptide that had been generated from either the cTfh CXCR3^+^, cTfh CXCR3^−^ or cNon-Tfh subset in 3 donors (**A**). For each clone, the peptide specificity, the amino acid sequence (AA), the CDR3 region of the ⍺ and β chains of the TCR and the family classification of J region (TRAJ and TRBJ) and V region (TRAV and TRBV) are shown. Clones are grouped based on the peptide recognised. The TCR ⍺ and β CDR3 region sequences of 75 HA-specific clones generated from the blood populations were determined, and the *ex vivo* V⍺ and Vβ repertoires of tonsil and blood Tfh and non-Tfh populations from the same subject were searched for matching sequences. Matches were detected only in donor 2. The frequency distributions of each of the individual clonotypes detected in the tonsil tTfh CXCR3^+^ (**B**) and peripheral blood cTfh CXCR3^+^ (**C**) repertoires are shown, with yellow circles indicating the sequences that matched to those of the HA-specific cTfh CXCR3^+^ clones generated from the peripheral blood of the same donor.

We next investigated whether the TCRα or TCRβ chain sequences identified in HA-specific T cell clones were detectable in the overall repertoire of CD4^+^ T cell subsets from both tonsil and peripheral blood, which had been evaluated by non-paired αβ TCR deep-sequencing. Since both the TCRα- and TCRβ-chain of each HA-specific T cell clone were sequenced, it was possible to determine the presence or the absence of each clonotype on the basis of detection of both chains in the subsets analyzed. Although only 75 HA-specific clones were sequenced, 3 cTfh CXCR3^+^ clones from donor 2 were found in the tTfh CXCR3^+^ repertoire from the same donor (**Figure 7B**) and two clones, also from donor 2, were detected in the cTfh CXCR3^+^ subset (**Figure 7C**). The tables show the frequency of the clonotype of interest observed in tonsil (**Figure S5B**) and in blood (**Figure S5C**) for both TCR chains. The presence of HA-specific Tfh clones initially derived from PBMCs in the tTfh CXCR3^+^ repertoire of the same donor confirmed the direct relationship between antigen-specific circulating and tonsillar Tfh cells.

## DISCUSSION

The work presented herein analyses the clonotypic overlap of Tfh cell subsets from peripheral blood and tonsillar cell populations. Repertoire sharing was selectively detected between blood and tonsillar populations with similar phenotypic characteristics, i.e. cTfh CXCR3^+^ and tTfh CXCR3^+^, cTfh CXCR3^−^ and tTfh GC/tTfh CXCR3^−^, and cNon-Tfh and tNon-Tfh subsets. These results suggest that, in a steady state condition, the clonal composition of the CXCR5^+^PD-1^+^ circulating cTfh population reflects that of tonsil-resident tTfh cells. In addition, the TCR repertoire analysis provided evidence of a clear distinction between Tfh and non-Tfh cell populations in both tissues. These observations were confirmed by analysis of antigen-specific T cell clones recognizing HA from influenza virus. Therefore, separation of Tfh cells from the total memory CD4^+^ T cell pool is necessary to investigate the specificity of the follicular T cell response.

In line with previous observations, effector Tfh GC (CXCR5^hi^PD-1^hi^) were the major sources of IL-21 and IL-4 and were only present in tonsil (Dan et al., 2019; He et al., 2013; Heit et al., 2017; Locci et al., 2013). The presence of Tfh GC cells in tonsils suggested a probable ongoing immune response in this tissue (Baumjohann et al., 2013; Hansen et al., 2016; Havenar-Daughton et al., 2016), which was not surprising considering that tonsillectomy is often a therapeutic surgical solution for recurrent chronic viral and bacterial infections (Dan et al., 2019; Grivel and Margolis, 2009; Rao, 2018). We also showed that Tfh GC cells are enriched in CXCR3^−^ cells and produced little IFN-γ (Ma et al., 2009; Ploquin et al., 2011). The lack of CXCR3 expression in Tfh GC cells is consistent with their specific effector function inside the GC, where they do not need to directly face inflammation but actively participate in the formation of neutralizing antibodies (Djuretic et al., 2007; He et al., 2013). Phenotypic features highlighted similarities, in particular between tTfh CXCR3^+^ and cTfh CXCR3^+^ subsets, but also heterogeneity between blood and tonsil populations (Chevalier et al., 2011; He et al., 2013). These differences may be driven by stimuli present in the respective local environments (Wong et al., 2016) and reflect selective recruitment, retention or differentiation of populations at particular sites (Durand et al., 2019).

To determine the relationship between circulating and tonsillar Tfh cell populations better, we considered a parameter that does not change with cellular location or differentiation, the TCR repertoire. Given the vast diversity of the TCR repertoire within each individual (10^8^), the number of cells sequenced (10^3^-10^5^) always constrains TCR repertoire studies and poses caveats requiring the use of technical or *in silico* replicates to confirm the primary observations. However, TCR deep-sequencing highlighted correlations between tonsil and blood Tfh and non-Tfh populations. Although RNA-based TCR sequencing provides relatively good repertoire coverage, it is not possible to calculate the absolute frequency of each clonotype with the populations analyzed (Heather et al., 2017; Rosati et al., 2017). However, we combined both weighted (the F2 metric) and unweighted (% of shared clonotypes) analysis and observed similar trends, validating the repertoire overlap observed between populations.

Effector Tfh GC cells from tonsil and tTfh CXCR3^−^ cells showed a moderate repertoire overlap with cTfh CXCR3^−^ cells from peripheral blood. Although a Tfh GC/cTfh CXCR3^−^ relationship between tonsil and blood has been previously documented in terms of gene expression (Locci et al., 2013), our analysis of donor-matched tissues directly demonstrates that the same Tfh clones are present in both tonsil and the blood CXCR5^+^PD-1^+^ cTfh compartment. The relatively modest overlap might be explained by the specific and localized role of Tfh GC cells during their effector phase in the GC (Groom and Luster, 2011). Consistent with this, we observed that shared clonotypes are more expanded in tonsil than in blood, suggesting selective retention of a locally-expanded Tfh population. The clonotype sharing between tTfh GC and tTfh CXCR3^−^ cells may reflect the fact that both are enriched for cells recognizing locally-presented antigens, with Tfh GC potentially originating from or sharing common precursors with Tfh CXCR3^−^ (Crotty, 2018; Locci et al., 2013). By contrast, the cTfh CXCR3^−^ population probably has a much more promiscuous origin, as it is likely to contain antigen-experienced Tfh cells that have exited from lymphoid tissues before or after becoming effector Tfh GC during both the ongoing and prior immune responses. It is also possible that cTfh cells are recruited from the periphery in response to local chronic infection in tonsils.

This study does not address the differentiation and the generation of Tfh CXCR3^+^ cells, but the convergence in clonotypic repertoire observed between peripheral blood and tonsil was striking. This suggests a high mobility of these types of cells from secondary lymphoid organs to the peripheral circulation or vice versa. The minor, but consistent, repertoire sharing between Tfh CXCR3^+^ and Tfh CXCR3^−^ cells in tonsil may result from tTfh cell differentiation from one phenotype to the other or plasticity among CXCR3^+^ and CXCR3^−^ phenotypes, previously observed in canonical T helper subsets (Becattini et al., 2014; Hegazy et al., 2010).

The repertoire sharing observed between non-Tfh cells in tonsil and blood confirms the communication between the two tissues and suggests that after priming CD4^+^ T cells from secondary lymphoid organs may exit into the peripheral circulation to pursue their effector function, but also that circulating non-Tfh cells may be recruited to the tonsil to react against ongoing chronic infection. Despite the consistent overlap, the weighted analysis showed that the size of clonotypes differed between the two compartments, indicating that these populations, which include multiple different subsets of CD4^+^ T cells, e.g. Th1, Th2 and Th17 cells, have a different clonotype distribution, which probably reflects differing retention and expansion in different locations.

In order to understand the magnitude of the observed repertoire overlaps better, it is necessary to consider the size of the samples analyzed (10^3^-10^5^) in comparison to the entire CD4^+^ T cell repertoire (10^8^) in the human body. Importantly, it was observed that when re-sampling exactly the same set of sequences *in silico* or by experimentally sequencing cells independently sorted from the same tissue sample, the repertoire overlap is never complete and it decreases with increasing population diversity. These quantitative results suggest that the degree of overlap observed experimentally between separately-sampled populations provides an underestimate of the considerable repertoire overlap between blood and tonsil Tfh populations.

Extending the analysis to antigen-specific populations, we showed using the T cell library technique (Geiger et al., 2009) that HA-specific CD4^+^ T cells were enriched in the cTfh CXCR3^+^ subset and the cNon-Tfh population, which contains the canonical CD4^+^ Th1 subsets. The number of HA-specific clones generated from the cTfh CXCR3^+^ and cNon-Tfh compartments was also much higher than that generated from the cTfh CXCR3^−^ pool. The CD4^+^CXCR3^+^ bias in the HA-specific response is in line with previous reports (Palladino et al., 1991; Pedersen et al., 2014; Valkenburg et al., 2018; Yang et al., 2013; Yoon et al., 2017; Zielinski et al., 2011). Notably, in one of the 3 donors studied, the TCR sequences of blood-derived HA-specific cTfh CXCR3^+^ clones were found in the tTfh CXCR3^+^ repertoire, supporting a common clonal origin of antigen-specific Tfh cells in tonsil and the circulation. These results from analysis of antigen-specific cells provide strong support for the conclusions drawn from TCR deep sequencing that the repertoires of cTfh and tTfh cells are very similar to each other but distinct from non-Tfh populations. Our observation of some repertoire sharing between Tfh GC and tTfh CXCR3^+^ populations is consistent with the hypothesis that some of the cells participating in a GC reaction originate from the tTfh CXCR3^+^ pool (Crotty, 2018; He et al., 2013). Whether a few Tfh GC cells, originally primed in a Th1 environment, are surviving to become memory Tfh Th1-like (or CXCR3^+^) cells in the lymphoid organs and later in the periphery is not clear (Crotty, 2018). Tfh GC cells may revert via a tTfh CXCR3^−^ stage to a tTfh CXCR3^+^ phenotype, as we observed some modest repertoire overlap between all three of these tTfh populations, as well as repertoire overlap unique to the tTfh CXCR3^−^ and tTfh CXCR3^+^ populations. These observations may apply to other Tfh cells with different phenotypes, similarly to canonical Th cell subsets, but future work focusing on different pathogens that drive Th2 or Th17 responses is needed to elucidate this further.

Our TCR repertoire analysis also suggested very little overlap between antigen-experienced Tfh and non-Tfh cell populations in blood and tonsil tissues under steady state conditions. However, our results do not exclude the possibility that memory CD4^+^ T cells, which are known to exhibit plasticity (Becattini et al., 2014; Caza and Landas, 2015; Hegazy et al., 2010; O’Shea et al., 2010), may alter their phenotype and become Tfh cells during an acute immune response. Analysis of the peptides recognized by HA-specific CD4^+^ T cell clones showed a wide diversity of specificities and a surprisingly small overlap in the peptides recognized by Tfh and non-Tfh subsets. The distinction between Tfh and non-Tfh subsets in peripheral blood was uniquely confirmed by the combination of TCR sequencing and peptide-specificity testing for cTfh CXCR3^+^, cTfh CXCR3^−^ and cNon-Tfh HA-reactive T cell clones, results from which indicated that these cells were generated from different naïve clones. Non-Tfh cells constitute the majority of the CD4^+^ T cell population in blood (PD-1^+^CXCR5^+^ cTfh comprise only 15% of memory and 6% of total blood CD4^+^ T cells) and, for many years, measurement of total CD4^+^ T cell responses in PBMCs has been used to assess the magnitude of helper T cell responses in the context of B cell and antibody responses. The findings reported here show that evaluation of the overall antigen-specific CD4^+^ T cell response will not give an accurate picture of the response in the Tfh compartment. Our results show that there is very little clonal overlap between *bona fide* helper Tfh cells, particularly Tfh GC, and CXCR5^−^CD4^+^ memory T cells in the blood or tonsil, and there were marked differences in the epitope specificity of HA-reactive cTfh and cNon-Tfh cells. However, there were significant overlaps between the CXCR5^+^ Tfh populations in blood and Tfh cells in lymphoid tissues. Therefore, in order to investigate the specificity of true Tfh responses, it will be necessary to identify circulating Tfh cells and to study their antigen reactivity. These findings have important implications for evaluating the breath and the specificity of immune responses in clinical trials of vaccines aimed at stimulating protective antibody responses.

## ACKNOWLEDGMENTS

We thank the donors for their willing participation in this study, the research nurses Deborah Barker and Dawn Young, James Ramsden FRCS and his surgical team at the John Radcliffe Hospital in Oxford, and Prof Stephen Harrison for provision of the haemagglutinin protein (Wisconsin strain). This work was supported by the National Institute of Allergy and Infectious Diseases at the National Institutes of Health (Centre for HIV/AIDS Vaccine Immunology-Immunogen Discovery, grant UM1 AI 00645), the Medical Research Council (MR/K012037), the Wellcome Trust (100326/Z/12/Z) and the Ministry of Education and Science of the Russian Federation (14.W03.31.0005). Pe.B., A.J.M., T.L. and S.C.G. are Jenner Institute Investigators. D.A.P. is a Wellcome Trust Senior Investigator.

## AUTHOR CONTRIBUTIONS

E.B. performed and designed the experiments, analyzed the data and wrote the first draft of the manuscript with supervision and support from S.L.C., Pe.B. and A.J.M; A.N.D. and M.M. performed TCR sequencing and data analysis with help and supervision from D.M.C; K.L. and J.E.M. performed TCR sequencing and data analysis with help and supervision from D.A.P; T.L. and S.C.G. set up and obtained ethical approval for the clinical study; Pa.B. performed statistical analysis and assisted E.B. with data analysis; Pe.B. and A.J.M. directed and supervised the project and edited the manuscript. All authors contributed intellectually and read and approved the final version of the manuscript.

## METHODS

### Experimental model and subject details

#### Human studies

Volunteer adult donors (16–50 years old) admitted for elective tonsillectomy at the John Radcliffe Hospital in Oxford were consented to donate peripheral blood (80–100mL) on the day of the surgery (**Table 1**). Patients were excluded from the study if they were affected by immunodeficiency, serious infection (e.g. Hepatitis B, Hepatitis C or HIV infection), lymphoid malignancy, cytogenetic disorders, history of cancer, patients with any pre-existing heart disease or blood clotting disorder. The existing cohort LABFLU002 was amended and ethically approved by the REC Oxford B Committee (13/SC/0152) and informed consent was obtained from all subjects. Peripheral blood samples for T cell library interrogations were collected from healthy donors via the UK National Blood Service.

**Table 1.** The table lists the donor ID, age, gender and indication for surgery of the donors from which tonsil and blood samples have been collected.

| Donor ID | Age | Gender | Indication for surgery |
| --- | --- | --- | --- |
| 1 | 42 | M | Sleep apnoea |
| 2 | 22 | F | Chronic recurrent tonsillar infection |
| 3 | 19 | M | Chronic recurrent tonsillar infection |
| 4 | 21 | M | Chronic recurrent tonsillar infection |
| 5 | 41 | M | Chronic recurrent tonsillar infection |
| 6 | 42 | M | Quinsy |
| 7 | 23 | F | Chronic recurrent tonsillar infection |
| 8 | 26 | M | Chronic recurrent tonsillar infection |
| 9 | 25 | F | Chronic recurrent tonsillar infection |
| 10 | 35 | F | Quincy and chronic recurrent tonsillar infection |
| 11 | 23 | F | Chronic recurrent tonsillar infection |
| 12 | 23 | F | Quincy and chronic recurrent tonsillar infection |
| 13 | 24 | F | Chronic recurrent tonsillar infection |
| 14 | 36 | F | Chronic recurrent tonsillar infection |
| 15 | 23 | M | Chronic recurrent tonsillar infection |
| 16 | 21 | F | Chronic recurrent tonsillar infection |

### Method details

#### Human sample processing

Tonsil nodules from adult donors were collected within 4 hours of surgery into RPMI glutamine [-] (Invitrogen) supplemented with non-essential amino acids (1%, Invitrogen), sodium pyruvate (1%, Invitrogen), glutamine (1%, Invitrogen), pooled AB human sera (5%, UK National Blood Service), β-mercaptoethanol (0.1%, Invitrogen) and penicillin/streptomycin (1%, Invitrogen). All samples were kept on ice during transportation. Tonsillar tissues were cleared from fatty and necrotic areas and cut into sections at a thickness of 2–3 mm using scalpels and scissors. The sections were then passed through a 70μm cell strainer and filtered through a 40μm cell strainer (Grivel and Margolis, 2009). Manually processed tonsils and peripheral blood samples were diluted 1:2 with Hank’s Balance Salt Solution (Sigma-Aldrich) and layered over 20 mL of Histopaque-1077 (Sigma-Aldrich). Mononuclear cells were isolated via density gradient centrifugation (800 g for 30 min). Flow cytometric analyses of phenotypic markers were performed directly *ex vivo*. CD14^+^ cells were enriched from PBMCs via magnetic separation using CD14 MicroBeads (Miltenyi Biotec).

#### Phenotypic analysis and intracellular cytokine staining

Mononuclear cells from tonsil and blood samples were washed in Dulbecco’s PBS (Sigma-Aldrich). For phenotypic analyses, cells were labelled with Live/Dead Fixable Aqua and surface stained with anti-CD4–APC-Cy7, anti-CD45RA–PE-TxRed, anti-CXCR3–PE-Cy5, anti-CXCR5–AF488, anti-ICOS–BV711, anti-PD-1–BV421 and the dump markers anti-CD8, anti-CD14, anti-CD16, anti-CD19, anti-CD25 and anti-CD56 (all conjugated to PE-Cy7) for 15 min at 37°C (reagent details in **Table 2**). For cytokine production, cells were stimulated with phorbol 12-myristate 13-acetate (2×10^−7^ M, Sigma-Aldrich) and ionomycin (1 μg/mL, Sigma-Aldrich) for 5 hr at 37°C. Brefeldin A was added after 2 hr (10μg/mL, Sigma-Aldrich). Cells were then surface stained as described above, fixed/permeabilized with Cytofix/Cytoperm 1X Solution (BD Pharmingen), and stained in Permwash 1X Solution (BD Pharmingen) with anti-CD4–APC-Cy7, anti-IFN-γ–AF700, anti-IL-4–PE and anti-IL-21–AF647 for 20 min on ice (reagent details in **Table 1**). The flow cytometry panel was internally validated using individually stained CompBeads (BD Biosciences). Stained samples were acquired using an LSR Fortessa (BD Biosciences) and analyzed using FlowJo v10.3 (Tree Star).

**Table 2.** Flow cytometry antibody list.

| Marker | Fluorochrome | Supplier | Clone | Dilution |
| --- | --- | --- | --- | --- |
| Live/Dead Fixable Aqua | Aqua | Invitrogen | L34957 | 1:1000 |
| Anti-CD3 | Brilliant Violet 605 | Becton Dickinson | UCHT1 | 1:50 |
| Anti-CD4 | APC-Cy7 | Becton Dickinson | RPA-T4 | 1:40 |
| Anti-CD45RA | PE-TxRed | Invitrogen | MEM-56 | 1:100 |
| Anti-CXCR3 | PE-Cy5 | Becton Dickinson | 1C6/CXCR3 | 1:20 |
| Anti-CXCR5 | Alexa Fluor 488 | Becton Dickinson | RF8B2 | 1:20 |
| Anti-ICOS | Brilliant Violet 711 | Becton Dickinson | DX29 | 1:50 |
| Anti-ICOS | PE | BioLegend | C398.4A | 1:40 |
| Anti-PD-1 | Brilliant Violet 421 | Becton Dickinson | EH12.1 | 1:40 |
| Anti-CD8 | PE-Cy7 | Becton Dickinson | SK1 | 1:50 |
| Anti-CD14 | FITC | Becton Dickinson | M5E2 | 1:50 |
| Anti-CD14 | PE-Cy7 | Becton Dickinson | M5E2 | 1:50 |
| Anti-CD16 | PE-Cy7 | Becton Dickinson | 3G8 | 1:50 |
| Anti-CD19 | PE-Cy7 | Becton Dickinson | SJ25C1 | 1:50 |
| Anti-CD25 | PE-Cy7 | Becton Dickinson | 2A3 | 1:50 |
| Anti-CD56 | PE-Cy7 | Becton Dickinson | B159 | 1:50 |
| Anti-IFN-γ | Alexa Fluor 700 | Becton Dickinson | B27 | 1:150 |
| Anti-IL-4 | PE | Becton Dickinson | MP4-25D2 | 1:80 |
| Anti-IL-21 | Alexa Fluor 647 | Becton Dickinson | 3A3-N2.1 | 1:40 |

#### CD4^+^ T cell library

CD4^+^ and CD14^+^ cells were enriched from the PBMCs of 3 healthy donors via magnetic separation using the corresponding MicroBeads (Miltenyi Biotec). The CD14^+^ fraction was cryopreserved in liquid nitrogen. CD4^+^ T cells were stained with Live/Dead Fixable Aqua, anti-CD3–BV605, anti-CD4–APC-Cy7, anti-CD45RA–PE-TxRed, anti-CXCR3–PE-Cy5, anti-CXCR5–AF488, anti-PD-1–BV421 and the dump markers anti-CD8, anti-CD14, anti-CD16, anti-CD19, anti-CD25 and anti-CD56 (all conjugated to PE-Cy7) for 15 min at 37°C (reagent details in **Table 2**). Circulating live CD3^+^CD4^+^CD45RA^−^ cells were sorted into cTfh CXCR3^+^ (CXCR3^+^CXCR5^+^PD-1^+^), cTfh CXCR3^−^ (CXCR3^−^CXCR5^+^PD-1^+^) and cNon-Tfh (CXCR5^−^) subsets using a FACSAria III (BD Biosciences). Sorting was performed using a 4-way purity to obtain 3–5×10^5^ cells per subset at a minimum purity of 92%. Sorted CD4^+^ T cells were seeded at a limiting dilution of 1×10^4^ cells/mL in RPMI glutamine [-] (Invitrogen) supplemented with non-essential amino acids (1%, Invitrogen), sodium pyruvate (1%, Invitrogen), glutamine (1%, Invitrogen), pooled AB human sera (5%, UK National Blood Service), β-mercaptoethanol (0.1%, Invitrogen), penicillin/streptomycin (1%, Invitrogen) and IL-2 (500 U/mL, University of Oxford). Cells were expanded with 1μg/mL phytohemagglutinin (Rebel) in the presence of irradiated (45 Gy) allogeneic feeder cells from 3 different healthy blood donors (10^6^ feeder cells/mL). After 20 days, each line was screened for the capacity to proliferate in response to HA. Autologous CD14^+^ monocytes were irradiated at 45 Gy and incubated (3×10^5^ cells/mL) for 5 hr at 37°C with overlapping HA peptides (2 μg/mL) corresponding to Influenza A Wisconsin/67/2005 (BEI Research Resources Repository). Peptides were 13–17 amino acids long with a 7–12 residue overlap spanning the entire length of HA. Negative control wells incorporated dimethyl sulfoxide (0.045%, Sigma-Aldrich), and positive control wells incorporated IL-2 (500 U/mL, University of Oxford). An aliquot of 2.5×10^6^ cells/mL from each expanded CD4^+^ T cell line was added to the monocytes after washing and resting in fresh culture medium without IL-2 for 5 hr. After 3 days, 1 μCi/mL [^3^H]-thymidine was added to the cultures, and proliferation was measured after 16 hr using a MicroBeta2 Counter (Perkin Elmer) (Campion et al., 2014; Geiger et al., 2009; Lindestam Arlehamn et al., 2013; Mele et al., 2017).

#### Antigen-specific CD4^+^ T cell clones

Frozen CD14-depleted PBMCs were defrosted and rested overnight at 37°C in RPMI glutamine [-] (Invitrogen) supplemented with non-essential amino acids (1%, Invitrogen), sodium pyruvate (1%, Invitrogen), glutamine (1%, Invitrogen), pooled AB human sera (10%, UK National Blood Service), β-mercaptoethanol (0.1%, Invitrogen) and penicillin/streptomycin (1%, Invitrogen). CD4^+^ cells were enriched via magnetic separation using CD4 MicroBeads (Miltenyi Biotec) and surface stained as described above. Among live lymphocytes, circulating CD3^+^CD4^+^CD45RA^−^ cells were sorted into cTfh CXCR3^+^ (CXCR3^+^CXCR5^+^PD-1^+^), cTfh CXCR3^−^ (CXCR3^−^ CXCR5^+^PD-1^+^) and cNon-Tfh (CXCR5^−^) subsets using a FACS Aria III (BD Biosciences). Sorted subsets (3×10^6^ cells/mL) were stimulated at a 3:1 ratio with irradiated (45 Gy) autologous CD14^+^ monocytes loaded as described above with overlapping HA peptides (2 μg/mL) corresponding to Influenza A California/04/2009 (GenScript). Peptides were 14–18 amino acids long with a 12 residue overlap spanning the entire length of HA. After 7 days, cells were stained with Live/Dead Fixable Aqua, anti-CD3–BV605, anti-CD4–APC-Cy7, anti-CD25–PE-Cy7, anti-CD45RA–PE-TxRed, anti-ICOS–PE and the dump marker anti-CD14 (conjugated to FITC) for 15 min at 37°C (reagent details in **Table 1**). Activated CD4^+^ T cells (CD3^+^CD4^+^CD14^−^CD25^+^ICOS^+^) were sorted using a FACSAria III (BD Biosciences) and seeded at 0.4 cells/well into 384-well plates (Corning). Cells were expanded with 1μg/mL phytohemagglutinin (Rebel) in the presence of irradiated (45 Gy) allogeneic feeder cells from 3 different healthy blood donors (10^6^ feeder cells/mL) in RPMI glutamine [-] (Invitrogen) supplemented with non-essential amino acids (1%, Invitrogen), sodium pyruvate (1%, Invitrogen), glutamine (1%, Invitrogen), pooled AB human sera (10%, UK National Blood Service), β-mercaptoethanol (0.1%, Invitrogen), penicillin/streptomycin (1%, Invitrogen) and IL-2 (500 U/mL, University of Oxford). After 14 days, T cell clones were identified and transferred into 96-well round-bottom plates (Corning). To confirm specificity, an aliquot of each clone (1×10^6^ cells/mL) was stimulated with the relevant overlapping peptide pool (2 μg/mL) after washing and resting in fresh culture medium without IL-2 for 5 hr. After 3 days, 1 μCi/mL [^3^H]-thymidine was added to the cultures, and proliferation was measured after 16 hr using a MicroBeta2 Counter (Perkin Elmer). Positive responses were defined as >1,000 cpm with a stimulation index >5 after background subtraction (cpm in wells lacking peptide). Clones were excluded from the analysis if the background count was >1,000 cpm.

#### Peptide-mapping of antigen-specific CD4^+^ T cell clones

Antigen-specific CD4^+^ T cell clones were mapped using a matrix of overlapping HA peptides corresponding to Influenza A California/04/2009, designed using “Deconvolute This!” (Precopio et al., 2008). The matrix consisted of 12 pools with 18 peptides per pool and a coverage of 3. An aliquot of each clone (1.5×10^6^ cells/mL) was added to the peptide matrix (2 μg/mL) after washing and resting in fresh culture medium without IL-2 for 5 hr. After 3 days, 1 μCi/mL [^3^H]-thymidine was added to the cultures, and proliferation was measured after 16 hr using a MicroBeta2 Counter (Perkin Elmer). Positive responses were defined as >1,000cpm with a stimulation index >5 after background subtraction (cpm in wells lacking peptide). Peptide specificity was confirmed similarly using single peptides identified from the matrix responses.

#### TCR deep-sequencing

Mononuclear cells from matched tonsil and peripheral blood samples were defrosted and rested overnight at 37°C in RPMI glutamine [-] (Invitrogen) supplemented with non-essential amino acids (1%, Invitrogen), sodium pyruvate (1%, Invitrogen), glutamine (1%, Invitrogen), pooled AB human sera (10%, UK National Blood Service), β-mercaptoethanol (0.1%, Invitrogen) and penicillin/streptomycin (1%, Invitrogen). Cells were then surface stained as described above. Among live CD3^+^CD4^+^CD45RA^−^ lymphocytes, circulating cells were sorted into cTfh CXCR3^+^ (CXCR3^+^CXCR5^+^PD-1^+^), cTfh CXCR3^−^ (CXCR3^−^CXCR5^+^PD-1^+^) and cNon-Tfh (CXCR5^−^) subsets, and cells from tonsils were sorted into tTfh GC (CXCR5^hi^PD-1^hi^), tTfh CXCR3^+^ (CXCR3^+^CXCR5^int^PD-1^int^), tTfh CXCR3^−^ (CXCR3^−^CXCR5^int^PD-1^int^) and tNon-Tfh (CXCR5^−^PD-1^−^) subsets. A maximum of 1.2×10^5^ cells per subset was sorted using a FACSAria III (BD Biosciences) into RLT buffer (Qiagen) containing 10% β-mercaptoethanol (Sigma). Sorted cells were stored at –80°C. Each subset from donors 5, 6 and 7 was sorted into 2 vials for technical replicates (**Table 3**). Total RNA was isolated using an RNeasy Mini kit (Qiagen). Unique molecular identifier (UMI)-labeled 5’-RACE TCRβ cDNA libraries were prepared using a Human TCR Profiling Kit (MiLaboratory LLC). All extracted RNA was used for cDNA synthesis, and all synthesized cDNA was used for PCR amplification. Libraries were prepared in parallel using the same number of PCR cycles and sequenced in parallel using a 150+150 bp NextSeq (Illumina).

**Table 3.** Number of cells sorted per subset and percentage of CD4^+^CD45RA^−^ (n.d.= non detected).

| N° cells sequenced and % CD4<sup>+</sup>CD45RA<sup>-</sup> | Peripheral blood |  |  | Tonsil |  |  |  |
| --- | --- | --- | --- | --- | --- | --- | --- |
| DONOR ID | Tfh CXCR3<sup>+</sup> | Tfh CXCR3<sup>-</sup> | Non-Tfh | Tfh GC | Tfh CXCR3<sup>+</sup> | Tfh CXCR3<sup>-</sup> | Non-Tfh |
| 1 | 23,000 | 50,000 | 50,000 | 100,000 | 100,000 | 100,000 | 100,000 |
|  | (2.7%) | (5.0%) | (57.3%) | (14.0%) | (4.6%) | (29.1%) | (15.8%) |
| 2 | 13,000 | 20,000 | 50,000 | 100,000 | 100,000 | 100,000 | 100,000 |
|  | (1.3%) | (2.0%) | (62.8%) | (4.4%) | (5.0%) | (21.3%) | (27.7%) |
| 3 | 8,000 | 17,000 | 80,000 | 123,000 | 119,000 | 120,000 | 120,000 |
|  | (0.7%) | (3.6%) | (84.0%) | (17.6%) | (7.5%) | (21.2%) | (21.4%) |
| 4 | 12,000 | 30,000 | 50,000 | 100,000 | 100,000 | 100,000 | 100,000 |
|  | (0.6%) | (1.6%) | (78.1%) | (5.8%) | (2.7%) | (22.2%) | (38.3%) |
| DONOR ID | Tfh CXCR3<sup>+</sup> | Tfh CXCR3<sup>-</sup> | Non-Tfh | Tfh GC | Tfh CXCR3<sup>+</sup> | Tfh CXCR3<sup>-</sup> | Non-Tfh |
| 5 (vial 1) | 4,000 | n.d. | 100,000 | 100,000 | 100,000 | 100,000 | n.d. |
|  | (0.5%) | n.d. | (79.5%) | (4.28%) | (3.7%) | (16.2%) | n.d. |
| 5 (vial 2) | 3,000 | n.d. | 100,000 | 100,000 | 100,000 | 100,000 | 100,000 |
|  | (0.5%) | n.d. | (79.5%) | (4.28%) | (3.7%) | (16.2%) | (26.7%) |
| 6 (vial 1) | n.d. | n.d. | 50,000 | 50,000 | 80,000 | 100,000 | 100,000 |
|  | n.d. | n.d. | (55.6%) | (5.5%) | (4.6%) | (18.5%) | (15.8%) |
| 6 (vial 2) | n.d. | n.d. | 50,000 | 50,000 | 100,000 | 100,000 | 100,000 |
|  | n.d. | n.d. | (55.6%) | (5.5%) | (4.6%) | (18.5%) | (15.8%) |
| 7 (vial 1) | n.d. | n.d. | n.d. | 100,000 | n.d. | n.d. | 100,000 |
|  | n.d. | n.d. | n.d. | (9.15%) | n.d. | n.d. | (23.8%) |
| 7 (vial 2) | n.d. | n.d. | n.d. | 100,000 | n.d. | 70,000 | 100,000 |
|  | n.d. | n.d. | n.d. | (9.15%) | n.d. | (12.4%) | (23.8%) |

#### TCR sequencing of antigen-specific CD4^+^ T cell clones

Antigen-specific CD4^+^ T cell clones were stained as described above for viability and surface expression of CD3 and CD4 (**Table 1**). A maximum of 5×10^3^ live CD3^+^CD4^+^ T cells was sorted into 100 μL of RNAlater (Ambion) using a FACSAria III (BD Biosciences). Sorted cells were stored at −80°C. All expressed *TRA* and *TRB* gene transcripts were amplified using an unbiased template-switch anchored RT-PCR (Quigley et al., 2011). Amplicons were subcloned, sampled and sequenced as described previously (Price et al., 2005). Gene use was assigned using the ImMunoGeneTics (IMGT) nomenclature (Lefranc et al., 2003).

#### Quantification of influenza-specific IgG

ELISA plates were coated with a commercial Trivalent Influenza Vaccine (0.75 μg/mL, Sanofi Pasteur) or HA protein from Influenza A/California/2009 (2μg/mL, BEI Research Resources Repository). After overnight incubation at 4°C, plates were washed 6 times with PBS containing 0.05% Tween-20, blocked with casein (Pierce) for 2 hr at room temperature, and washed again with PBS containing 0.05% Tween-20. Diluted plasma samples (1:500, 1:1,000, 1:2,000 and 1:4,000) were then added in duplicate. After incubation for 2 hr at room temperature, plates were washed again 6 times with PBS containing 0.05% Tween-20, incubated with alkaline phosphatase-conjugated goat anti-human IgG (1 μg/mL, Sigma-Aldrich) for 1 hr at room temperature, and washed again with PBS containing 0.05% Tween-20. Para-nitrophenylphosphate (Sigma-Aldrich) dissolved at a concentration of 1 mg/mL in diethanolamine buffer (Pierce) was then added for 10 min, and absorbance was read at 405 nm using a Microplate Reader (BioTek) with Open Gen5 software (BioTek). Results were plotted as arbitrary units and standardized against a pool of 4 plasma samples (Miura et al., 2008).

### Statistical analysis

#### Flow cytometry data

A stop gate was set during acquisition to ensure that the same number of events was acquired and for each sample. Gates were set on the data to determine the frequencies of the different subsets. Analogous subsets were compared between tonsil and peripheral blood using the Wilcoxon matched pairs test. Non-analogous subsets were compared within tonsil and peripheral blood using the Kruskal-Wallis test. *P<0.05, **P<0.01, ***P<0.001, ****P<0.0001. Boxes show means and quartiles, and whiskers show minima and maxima.

#### TCR deep-sequencing analysis

A total of 93×10^6^ TCRβ sequencing reads (up to 8×10^6^ reads per library) were obtained, from which 1.55×10^6^ unique UMI-labeled TCRβ cDNA molecules (up to 1.5×10^5^ molecules per library) were assembled using MIGEC (Shugay et al., 2014), using a threshold of at least 4 sequencing reads per UMI. In-frame CDR3β repertoires were extracted using MiXCR software (Bolotin et al., 2015). Each library contained from 500 to 3.5×10^4^ functional (in-frame, no stop-codons) CDR3β nucleotide clonotypes. Averaged TCR repertoire characteristics weighted by clonotype size (normalized Shannon-Wiener index) were calculated using VDJtools software (Shugay et al., 2015). As the number of cells in the populations sorted from different donors was variable, the frequency of each clonotype was normalized before analysis to allow an unbiased comparison between diverse subsets. Network analysis was performed using Cytoscape (https://cytoscape.org). The F2 metric for overlap between two populations was calculated as the sum of the geometric means between the frequencies of shared clonotypes, considering the top2000 clonotypes for each subset. This value was then standardized with the two maximum F2 metrics calculated for the top2000 clonotypes for each of the two populations. F2 metric, % shared clonotypes and diversity indices were analysed with non-parametric Kruskal-Wallis and Dunn’s multiple comparison tests. For each test: *P<0.05, **P<0.01, ***P<0.001, ****P<0.0001. Linear regression correlations were calculated using the square of the Pearson correlation coefficient (R^2^), the slope and the P value of the significance of the difference in the slope from zero (ns= not significant).

#### Bootstrap method

The complete set of clonotypes was randomly resampled with replacement to obtain two *in silico* replicates of each set of sequences. Using this method, the replicates and the original set had similar distributions, but likely different absolute numbers, for each clonotype. All downstream analyses were performed using the two resampled sets. Linear regression correlations between *in silico* replicates and diversity were calculated using the P value of the significance of the difference in the slope from zero.

#### T cell library analysis

Positive responses were defined as >3,000 counts per minute (cpm) with a stimulation index >5 after background subtraction (cpm in the negative control wells). Because CD4^+^ T cells were seeded at limiting dilution, it was possible to calculate the precursor frequency of responding cells per million according to the Poisson distribution (Campion et al., 2014; Geiger et al., 2009; Lindestam Arlehamn et al., 2013; Mele et al., 2017). The 95% confidence intervals were determined according to the modified Wald method (Harrell, 2013).

#### Antigen-specific CD4^+^ T cell frequencies

The frequencies of expanded antigen-specific CD4^+^ T cell clones were calculated based on the number of clones with confirmed specificity after expansion. Significantly diverse enrichments were identified using Kruskal-Wallis and Dunn’s multiple comparison tests were performed. For each test: *P<0.05, **P<0.01, ***P<0.001, ****P<0.0001.

