## Supplemental Figures for "CD4^+^ T follicular helper (Tfh) cells in human tonsil and blood are clonally convergent, but divergent from non-Tfh CD4^+^ cells"

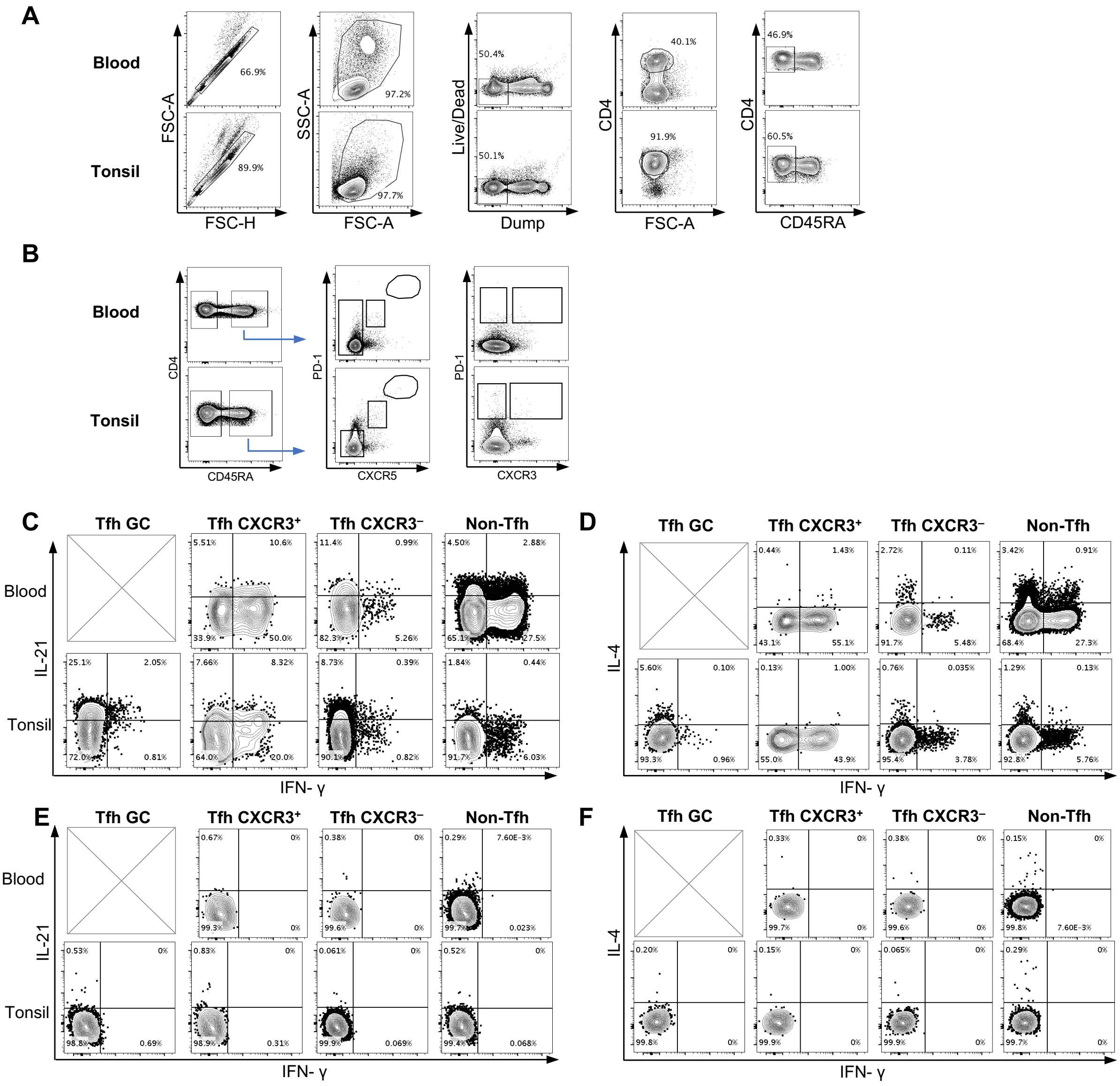
A
Blood
Tonsil
FSC-A
FSC-H
SSC-A
Live/Dead
Dump
CD4
CD45RA
B
PD-1
CXCR5
CXCR3
C
D
E
F
Tfh GC
Tfh CXCR3⁺
Tfh CXCR3⁻
Non-Tfh
IL-21
IL-4
IFN- γ

**Supplementary Figure 1. Gating strategy and cytokine production controls.**
Representative example of data from a matched tonsil and blood sample. From singlet cells (**A**), identified with Forward Scatter-Area (FSC-A) and Forward Scatter-Height (FSC-H), lymphocytes were gated based on FSC-A and Side Scatter-Area (SSC-A). The cells were then negatively gated for dead/dying cells and dump markers (CD16, CD56, CD8, CD25, CD14), then gated on the live $CD4^+$ population and $CD45RA^-$. Gates to define $CD4^+$ memory T cells populations were set based on the expression of PD-1, CXCR5 and CXCR3 in the naïve-enriched $CD4^+CD45RA^+$ T cell compartment (**B**).
Examples of IL-21, IFN-γ (**C**) and IL-4 (**D**) production as assessed by intracellular cytokine staining after PMA/ionomycin stimulation are shown for each $CD4^+CD45RA^-$ memory T cell subset considered. The gating was set based on the IL-21, IFN-γ (**E**) and IL-4 (**F**) staining of the unstimulated control for the same subset.

**A**

| Subset | Donor ID | N. shared clonotypes | $R^2$ | P value |
|---|---|---|---|---|
| cTfh CXCR3+ | 1 | 1012 | 0.94 | <0.0001 |
| | 2 | 1158 | 0.73 | <0.0001 |
| | 3 | 1499 | 0.7 | <0.0001 |
| | 4 | 1308 | 0.66 | <0.0001 |
| cTfh CXCR3– | 1 | 896 | 0.84 | <0.0001 |
| | 2 | 931 | 0.5 | <0.0001 |
| | 3 | 974 | 0.65 | <0.0001 |
| | 4 | 921 | 0.61 | <0.0001 |
| cNon-Tfh | 1 | 995 | 0.99 | <0.0001 |
| | 2 | 992 | 0.96 | <0.0001 |
| | 3 | 1110 | 0.88 | <0.0001 |
| | 4 | 891 | 0.82 | <0.0001 |
| tTfh GC | 1 | 1365 | 0.97 | <0.0001 |
| | 2 | 1452 | 0.98 | <0.0001 |
| | 3 | 1199 | 0.98 | <0.0001 |
| | 4 | 1412 | 0.94 | <0.0001 |
| tTfh CXCR3+ | 1 | 1275 | 0.99 | <0.0001 |
| | 2 | 1183 | 0.97 | <0.0001 |
| | 3 | 1026 | 0.99 | <0.0001 |
| | 4 | 1108 | 0.85 | 2.4 |
| tTfh CXCR3– | 1 | 965 | 0.78 | <0.0001 |
| | 2 | 964 | 0.77 | <0.0001 |
| | 3 | 790 | 0.56 | <0.0001 |
| | 4 | 802 | 0.5 | <0.0001 |
| tNon-Tfh | 1 | 875 | 0.76 | <0.0001 |
| | 2 | 788 | 0.73 | <0.0001 |
| | 3 | 748 | 0.73 | <0.0001 |
| | 4 | 600 | 0.59 | <0.0001 |

**B**

**Non-Tfh cell replicates**

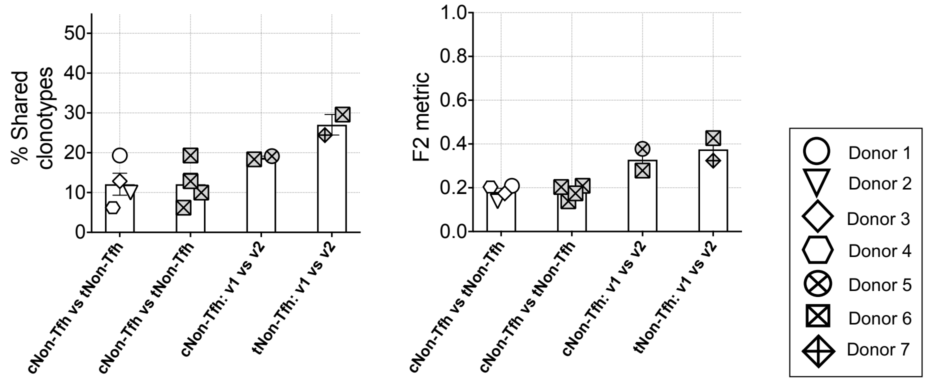


**C**

**Tfh cell replicates**

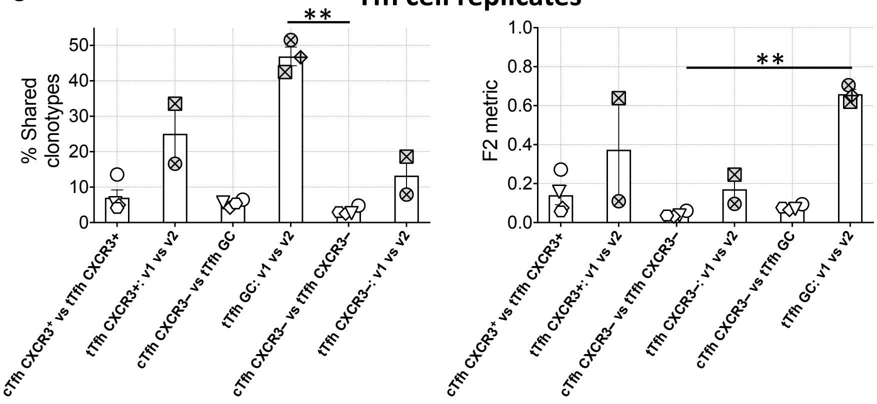

**Supplementary Figure 2. *In silico* resampling and experimental replicates analysis of the TCR Vβ CDR3 repertoire of blood and tonsil.** The clonotypes of each subset were resampled using the bootstrap method. The table (**A**) shows for each subset the number of shared clonotypes, the square of the Pearson correlation coefficient ($R^2$), and the P value of the significance of the difference in the slope from zero in the 4 donors.

From experimental replicates, the repertoire overlaps, percentage of shared clonotypes and normalized F2 metric (mean ± SEM), of the top2000 clonotypes of non-Tfh cells (**B**) from 2 different vials of 3 additional donors (donor 5, 6 and 7) were analyzed in comparison with the observed overlap from the first 4 donors (donor 1, 2, 3 and 4). Depending on the sample availability it was possible to calculate the overlap between two vials of the same subsets (v1 vs v2) or between blood and tonsil (cNon-Tfh vs tNon-Tfh). The same repertoire overlaps and comparison were made for Tfh cells (**C**) for the subsets available.

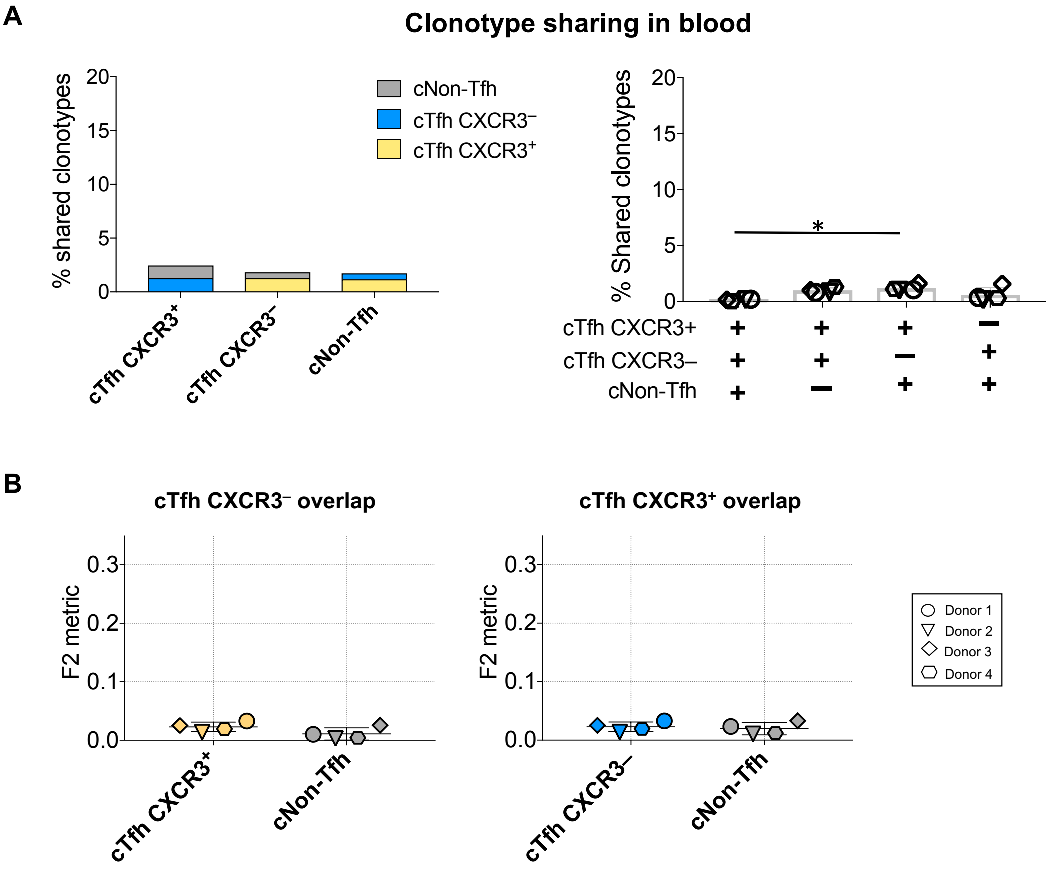


**C**

| Blood | cTfh CXCR3+ vs cTfh CXCR3– | | cTfh CXCR3+ vs cNon-Tfh | | cTfh CXCR3– vs cNon-Tfh | |
|---|---|---|---|---|---|---|
| Donor ID | Shared clonotypes | $R^2$ | Shared clonotypes | $R^2$ | Shared clonotypes | $R^2$ |
| **1** | 20 | 0.94 | 25 | 0.99 | 11 | 0.96 |
| **2** | 20 | 0.60 | 23 | 0.23 | 6 | 0.87 |
| **3** | 23 | 0.97 | 35 | 0.93 | 34 | 0.92 |
| **4** | 26 | 0.84 | 22 | 0.83 | 6 | 0.58 |

**Supplementary Figure 3. Tfh cell subsets from peripheral blood have a distinct repertoire from that of non-Tfh cells.** To compare the overlap between different populations of blood cells, the percentage of total shared clonotypes in addition to the uniquely shared clonotypes was calculated for each overlap analysis in the 4 donors (mean ± SEM) (**A**). The normalized F2 metric of the top2000 frequencies is also shown for cTfh $CXCR3^+$ and cTfh $CXCR3^-$ (**B**).

The table shows the number of shared clonotypes and the square of the Pearson correlation coefficient ($R^2$) (**C**).

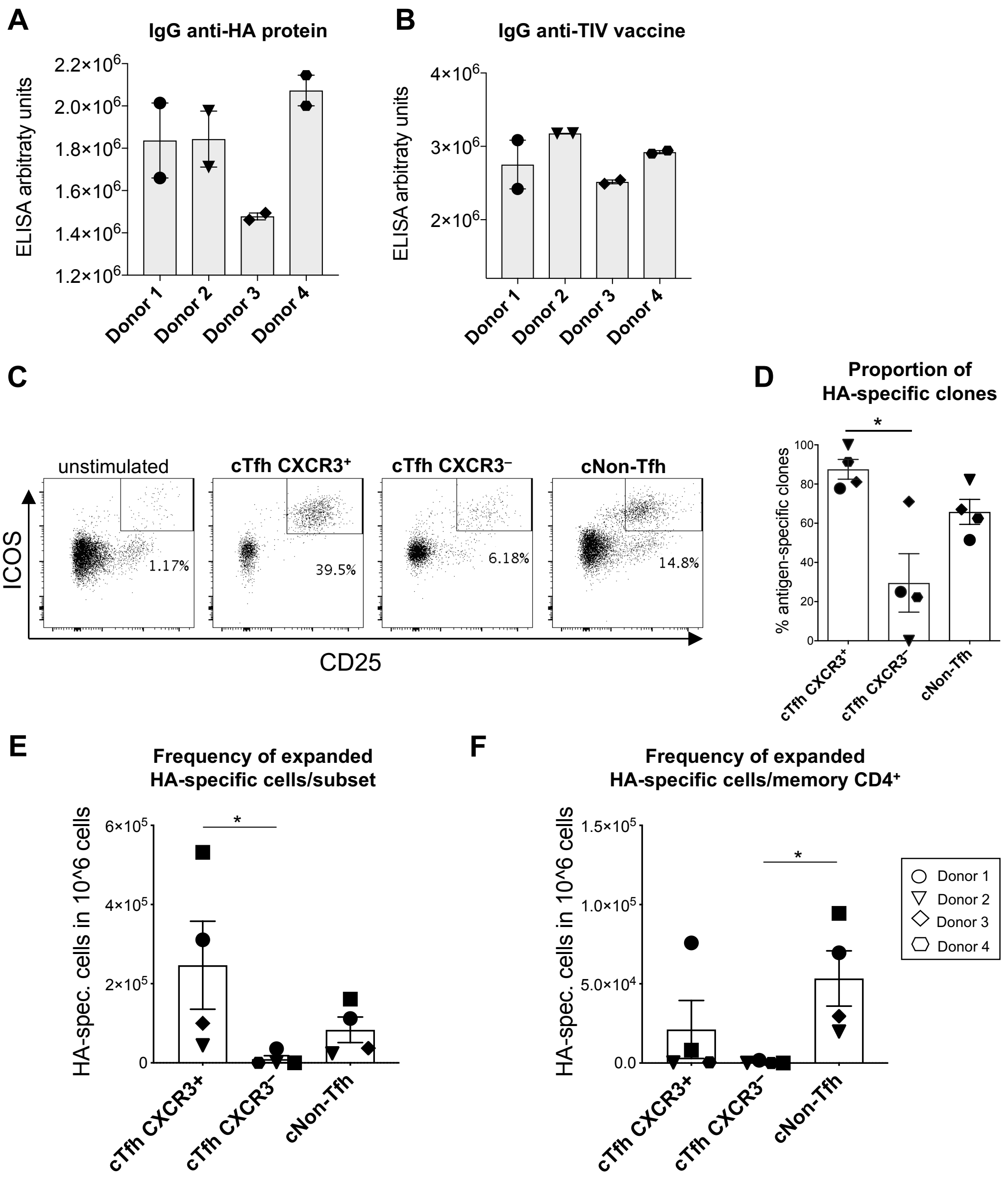

A
IgG anti-HA protein
ELISA arbitraty units
Donor 1
Donor 2
Donor 3
Donor 4
B
IgG anti-TIV vaccine
ELISA arbitraty units
C
unstimulated
cTfh CXCR3+
cTfh CXCR3−
cNon-Tfh
ICOS
CD25
1.17%
39.5%
6.18%
14.8%
D
Proportion of HA-specific clones
% antigen-specific clones
E
Frequency of expanded HA-specific cells/subset
HA-spec. cells in 10^6 cells
F
Frequency of expanded HA-specific cells/memory CD4+
Donor 1
Donor 2
Donor 3
Donor 4

**Supplementary Figure 4. Influenza antibody analysis and generation of HA-specific CD4$^+$ T cell clones.** IgG reactive with HA/California (**A**) and TIV (Trivalent Influenza Vaccine, from 2017) (**B**) in plasma samples from the 4 donors was analyzed by ELISA and is expressed in ELISA arbitrary units. Data presented are results from duplicate wells, and have been background subtracted (mean ± SEM). To generate HA-specific T cell clones from these 4 donors, memory CD4$^+$ T cell subsets sorted from peripheral blood were co-cultured with autologous monocytes (CD14$^+$) and overlapping peptides from HA/California, and activated memory CD4$^+$ T cells identified on the basis of co-expression of ICOS and CD25 were isolated by cell sorting after 7 days, as shown in the examples (from donor 3) in (**C**). A negative control (unstimulated cells from the cTfh CXCR3$^+$ subset) is also shown. The bar graph (**D**) shows the percentage of clones generated that were confirmed to be HA-specific on re-screening after expansion. Estimation of the number of HA-specific cells per $10^6$ total cells within each subset (**E**) or within memory CD4$^+$ (**F**) after 7 days expansion is represented from 4 donors (mean ± SEM).

**A**

Clones specific for peptide 313-330: HPITIGKCPKYVKSTKLR

| DONOR ID | CLONE TYPE | CLONE ID | TRAV | AA alpha_CDR3 | TRAJ | TRBV | AA beta_CDR3 | TRBJ |
|---|---|---|---|---|---|---|---|---|
| Donor 1 | cTfh CXCR3+ | 21 | 26-1 | CIVRVGREQGGKLI | 23 | 2 | CAKQGTGYNEQF | 2-1 |
| | | | 29/DV5 | CAAPEGTYKYI | 40 | | | |
| Donor 1 | cTfh CXCR3– | 1 | 13-1 | CAARTGAQKLV | 54 | 11-2 | CASTRTSGGANTGELF | 2-2 |
| Donor 1 | cTfh CXCR3– | 2 | 25 | CAGRGPNSNSGYALN | 41 | 18 | CASSQGYEQY | 2-7 |

**B**

| CLONE TYPE | CLONE ID | AA α CDR3 | TRAV | AA β CDR3 | TRBV | Vα frequency | Vβ frequency |
|---|---|---|---|---|---|---|---|
| cTfh $CXCR3^+$ | 21 | CAVRVGTGRRALT | 21 | CASSAGQATTGEQY | 5-1 | 7.42E-05 | 1.17E-04 |
| cTfh $CXCR3^+$ | 44 | CALKTGANNLF | 24 | CSAKAPGATQY | 20-1 | 2.47E-05 | 3.91E-05 |
| cTfh $CXCR3^+$ | 48 | CAVLLFMDSNYQLI | 22 | CAISEGGGSYGRKNIQY | 10-3 | 2.47E-05 | 2.61E-05 |

**C**

| CLONE TYPE | CLONE ID | AA α CDR3 | TRAV | AA β CDR3 | TRBV | Vα frequency | Vβ frequency |
|---|---|---|---|---|---|---|---|
| cTfh $CXCR3^+$ | 21 | CAVRVGTGRRALT | 21 | CASSAGQATTGEQY | 5-1 | 1.35E-04 | 5.35E-04 |
| cTfh $CXCR3^+$ | 57 | CAVLISSGSARQLT | 22 | CASSSHPTGTYGRNTEAF | 12-3 | 6.75E-04 | 7.49E-04 |

**Supplementary Figure 5. TCR CDR3 sequences of additional HA-specific cTfh clones from donor 1 and CDR3 sequences of cTfh CXCR3$^+$ clones detected in the tTfh CXCR3$^+$ subset from tonsil and blood in donor 2.**

The table lists additional clones from donor 1, specific for HA H1/California peptide 313-330 generated from cTfh CXCR3$^+$ and cTfh CXCR3$^-$ (**A**). For each clone, the CDR3 region of the α and β chains of the TCR and the family classification of the J region (TRAJ and TRBJ) and the V region (TRAV and TRBV) are shown.

The tables in **B** and **C** list the clones from donor 2 for which matching sequences were found in tonsil tTfh CXCR3$^+$ (**B**) and blood cTfh CXCR3$^+$ repertoires (**C**). The amino acid (AA) sequences of the CDR3 regions of the TCR α and β chains, family classification of the V regions (TRAV and TRBV) and the clonotype frequency within the relevant population are shown.
